# Aging is associated with functional and molecular changes in distinct hematopoietic stem cell subsets

**DOI:** 10.1101/2023.08.08.552444

**Authors:** Ece Somuncular, Julia Hauenstein, Tsu-Yi Su, Özge Dumral, Charlotte Gustafsson, Efthymios Tzortzis, Aurora Forlani, Anne-Sofie Johansson, Robert Månsson, Sidinh Luc

## Abstract

Age is a risk factor for hematologic malignancies. Attributes of the aging hematopoietic system include increased myelopoiesis, impaired adaptive immunity, and a functional decline of the hematopoietic stem cells (HSCs) that maintain hematopoiesis. Changes in the composition of diverse HSC subsets have been suggested to be responsible for age-related alterations, however, the underlying regulatory mechanisms are incompletely understood in the context of HSC heterogeneity. In this study, we investigated how distinct HSC subsets, separated by CD49b, functionally and molecularly change their behavior with age. We demonstrate that blood lineage differentiation progressively shifts to a higher myeloid cellular output in both lymphoid-biased and myeloid-biased HSC subsets during aging. In parallel, we show that HSCs selectively undergo age-dependent gene expression and gene regulatory molecular changes in a progressive manner, which is initiated already in the pre-adult stage. Overall, our studies suggest that aging intrinsically alters both cellular and molecular properties of HSCs.

**Highlights:**

- With age a gradual shift towards myeloid differentiation occurs in both myeloid-biased and lymphoid-biased enriched HSC subsets.
- Age-related molecular changes preferentially occur in HSCs.
- Functionally distinct HSC subsets with high transcriptional similarity can be distinguished on the epigenetic level.
- HSC aging is associated with a progressive increase in chromatin accessibility.

---

Aging of an organism is associated with physiological changes in all organ systems and a progressive functional decline. The age-related functional impairment leads to difficulties in maintaining homeostasis, especially during stress. Therefore, aging has many health implications and is one of the main risk factors of developing cancer^1–4^. Normal aging of the hematopoietic system is associated with decreased competence of the immune system, onset of anemia, and an increased risk of hematologic disorders including myeloid malignancies^1–3^. Throughout life, the hematopoietic system and its components are maintained and replenished by hematopoietic stem cells (HSCs)^5^. It has been suggested that aging features of the hematopoietic system are due to functional alterations in the capacity of HSCs to maintain homeostasis. Hematopoietic stem cells increase in both frequency and number with age. However, the regenerative capacity of old HSCs is reduced compared to their young counterparts, indicating that the diminished function is partly counterbalanced by increased HSC numbers to maintain homeostasis^1, 2, 6^.

It is well-recognized that the HSC compartment is functionally diverse containing not only lineage-balanced HSCs, but also myeloid-, platelet-, and lymphoid-biased HSC subsets that preferentially generate cells from specific blood lineages^7–11^. Studies have suggested that the blood cell composition changes with age, with myeloid cells becoming predominant and lymphoid cells declining in number. Different models have been proposed to underlie the myeloid skewing of the hematopoietic system. In the HSC clonal composition model, where lineage differentiation potential of individual HSCs remains unchanged, the aging-related myeloid predominance is attributed to an increase in platelet-and myeloid-biased HSCs, and a decrease in lymphoid-biased HSCs^6, 9, 12–15^. On the contrary, in the cell-intrinsic model, changes in the differentiation properties of HSCs result in diminished ability to generate lymphoid cells, leading to an accumulation of myeloid cells^16^. Further studies are needed to elucidate whether distinct HSC subsets undergo age-dependent intrinsic functional changes that explains the myeloid bias of an aging hematopoietic system^1, 2, 17^.

Aging is associated with extensive gene expression and gene regulatory changes, especially in HSC proliferation and differentiation associated genes. Comprehensive epigenome studies have shown that gene loci associated with differentiation are hypermethylated, while loci correlated with HSC function are hypomethylated and display an increase in activating histone marks in aged HSCs^16, 18–21^. To what degree these molecular differences reflect the changing composition of functionally different HSC subsets in aging has not been elucidated. Epigenetic characterization of highly enriched lineage-biased HSC subsets has thus far not been widely performed due to limitations in prospectively isolating functionally distinct HSCs. The epigenetic changes associated with lineage-biased HSCs in aging therefore remain largely unexplored.

We have previously used the integrin CD49b as a prospective marker to distinguish between functionally different subsets within the primitive Lineage^−^Sca-1^+^c-Kit^+^ (LSK) CD48^−^ CD34^−^CD150^hi^ HSC compartment^11, 12, 22, 23^. We demonstrated that the CD49b^−^ subset is highly enriched in cells with myeloid bias, while the CD49b^+^ fraction mainly showed lymphoid-biased features. Furthermore, we showed that CD49b^−^ and CD49b^+^ HSCs were transcriptionally similar but had distinct chromatin accessibility profiles, suggesting that functional differences between lineage-biased HSCs are epigenetically regulated^22^.

In the present study, we assessed the functional and molecular changes of distinct HSC subsets phenotypically separated by CD49b in the pre-adult (juvenile period before sexual maturity and adulthood)^24, 25^, adult, and old stages of development. We found that cell proliferation and cell cycle kinetics dynamically change, and that more myeloid cells are gradually generated *in vivo* with age, from both the CD49b^−^ myeloid-biased and CD49b^+^ lymphoid-biased HSC subsets. Molecular characterization revealed age-dependent transcriptional and epigenetic changes that preferentially occurred in HSCs. Our studies demonstrate that aging is associated with progressive functional and molecular changes in both CD49b^−^ and CD49b^+^ HSCs, including altered blood lineage output, gene expression, and remodeling of the chromatin landscape.

## Results

### CD49b expression in the HSC compartment is conserved in aging

With age, there is an increased number of total bone marrow (BM) cells, myeloid cells, and phenotypic HSCs^1^. Since CD49b can subfractionate HSCs into lineage-biased subsets^22^, we examined whether the frequency of CD49b^−^ and CD49b^+^ cells alter with age. Given that HSCs acquire an adult phenotype at 3-4 weeks after birth^26^, we investigated the phenotypic LSKCD48^−^ CD34^−^CD150^hi^ (CD150^hi^) HSC compartment in young (∼1 month old), adult (∼2-4 months old), and old (∼1.5-2 years old) mice. The CD150^hi^ fraction was significantly expanded in old mice, compared to young and adult mice (Fig. 1a and Extended Data Fig. 1a), consistent with previous reports^12^. CD49b subdivided CD150^hi^ cells into CD49b^−^ and CD49b^+^ subfractions with similar ratios in all age groups (Extended Data Fig. 1b,c). In line with an expanded CD150^hi^ fraction, the total frequency and number of phenotypic CD49b^−^ and CD49b^+^ cells significantly increased in old mice, compared to young and adult mice (Fig. 1b,c).

**Fig. 1.**
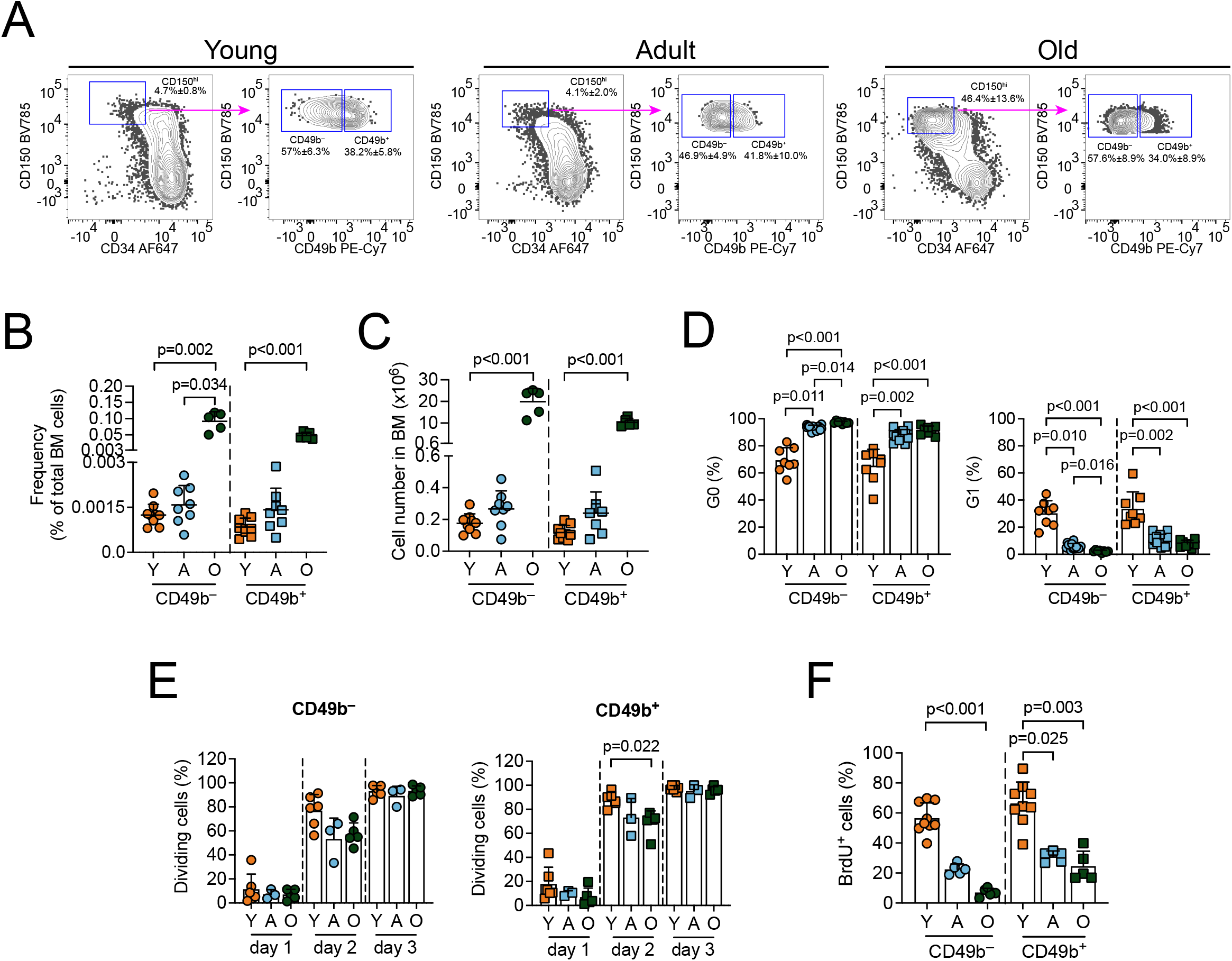
CD49b expression in the HSC compartment is conserved in aging. **a**, FACS profiles and gating strategy of the phenotypic HSC compartment (Lineage^−^Sca-1^+^c-Kit^+^ (LSK) CD48^−^ CD34^−^CD150^hi^), with further separation using CD49b, in young, adult, and old mice. Frequency of parent gates are shown. See Extended Data Fig. 1a for the full gating strategy. **b**, Total frequency of CD49b^−^ and CD49b^+^ HSC subsets in young (n = 9, 2 experiments), adult (n = 8, 5 experiments), and old (n = 5, 5 experiments) mice. **c**, Total numbers of CD49b^−^ and CD49b^+^ HSC subsets in young (n = 9, 2 experiments), adult (n = 8, 5 experiments), and old (n = 5, 5 experiments) mice. **d**, Frequency of CD49b^−^ and CD49b^+^ HSCs in G0 (left) and G1 (right) of young (n = 8, 2 experiments), adult (n = 15, 6 experiments), and old (n = 8, 5 experiments) mice. **e**, Frequency of cell divisions from cultured single cells of CD49b^−^ (left) and CD49b^+^ (right) HSCs at days 1-3 from young (n = 6 mice, 2 experiments, n_CD49b_^−^ = 297 cells, n_CD49b_^+^ = 214 cells), adult (n = 3 mice, 2 experiments, n_CD49b_^−^ = 112, n_CD49b_^+^ = 107), and old (n = 5 mice, 4 experiments, n_CD49b_^−^= 357, n_CD49b_^+^ = 381) mice. **f**, Frequency of BrdU^+^ CD49b^−^ and CD49b^+^ HSCs from young (n = 9, 2 experiments), adult (n = 5, 1 experiment), and old (n = 5, 2 experiments) mice. Mean ± s.d. is shown. The statistical analysis was performed with Kruskal-Wallis with Dunn’s multiple comparison test. Y, young; A, adult; O, old. See also Extended Data Fig. 1.

Hematopoietic stem cells largely reside in a quiescent state. However, it was previously demonstrated that nearly all CD49b^−^ HSCs within the CD150^hi^ compartment were in G0, while a significantly higher frequency of CD49b^+^ HSCs were in G1^22^. To investigate cell cycle changes in CD49b^−^ and CD49b^+^ subsets throughout aging, we performed cell cycle analysis using Ki-67 staining in young, adult, and old mice (Extended Data Fig. 1d). Cells from both CD49b subsets became progressively more quiescent (G0 phase) with age, with a corresponding decreased number of cells in G1 (Fig. 1d). Furthermore, CD49b^−^ HSCs from adult and old mice were significantly more quiescent than their CD49b^+^ counterparts (Extended Data Fig. 1e). However, no difference was observed between the two CD49b subsets in young mice (Extended Data Fig. 1e). To further assess cycling differences, we analyzed the cell division kinetics of individual CD49b^−^ and CD49b^+^ cells during aging. Compatible with the high frequency of cells in G0, most cells had not divided on the first day (Fig. 1e). By the second day, more CD49b^−^ and CD49b^+^ cells from young mice had undergone cell division compared to their adult and old counterparts, consistent with the reduced frequency of young CD49b^−^ and CD49b^+^ cells in G0, however, the difference was only statistically significant for the CD49b^+^ subset (Fig. 1d,e). To examine the cell proliferation properties *in vivo*, we performed 5-bromo-2’-deoxyuridine (BrdU) incorporation assay (Extended Data Fig. 1f). BrdU labeling drastically decreased with age in both CD49b subsets (Fig. 1f). Furthermore, adult and old CD49b^+^ subsets proliferated more than their corresponding CD49b^−^ cells, while both CD49b subsets in young mice were equally proliferative (Extended Data Fig. 1g), consistent with cell cycle and cell division results. Altogether, our results show that the number of both CD49b^−^ and CD49b^+^ cells dramatically increases with age. Moreover, while both subsets progressively become more quiescent and less proliferative with age, CD49b^−^ is the more dormant subset.

### The lineage repopulation patterns of multipotent CD49b subsets change with age

The differentiation potential of HSCs has been suggested to alter with age^1, 6, 16^. To functionally assess the differentiation ability of CD49b^−^ and CD49b^+^ subsets throughout aging, we first investigated their lymphoid, myeloid, and megakaryocyte potential at the clonal level *in vitro*. In all age groups, both CD49b subsets generated B lymphocytes and myeloid cells (Fig. 2a). Similarly, CD49b^−^ and CD49b^+^ cells from all ages efficiently generated megakaryocytes with no significant differences (Fig. 2b). Thus, although the frequency and number of CD49b^−^ and CD49b^+^ cells dramatically increased, the lineage differentiation ability *in vitro* was preserved with age.

**Fig. 2.**
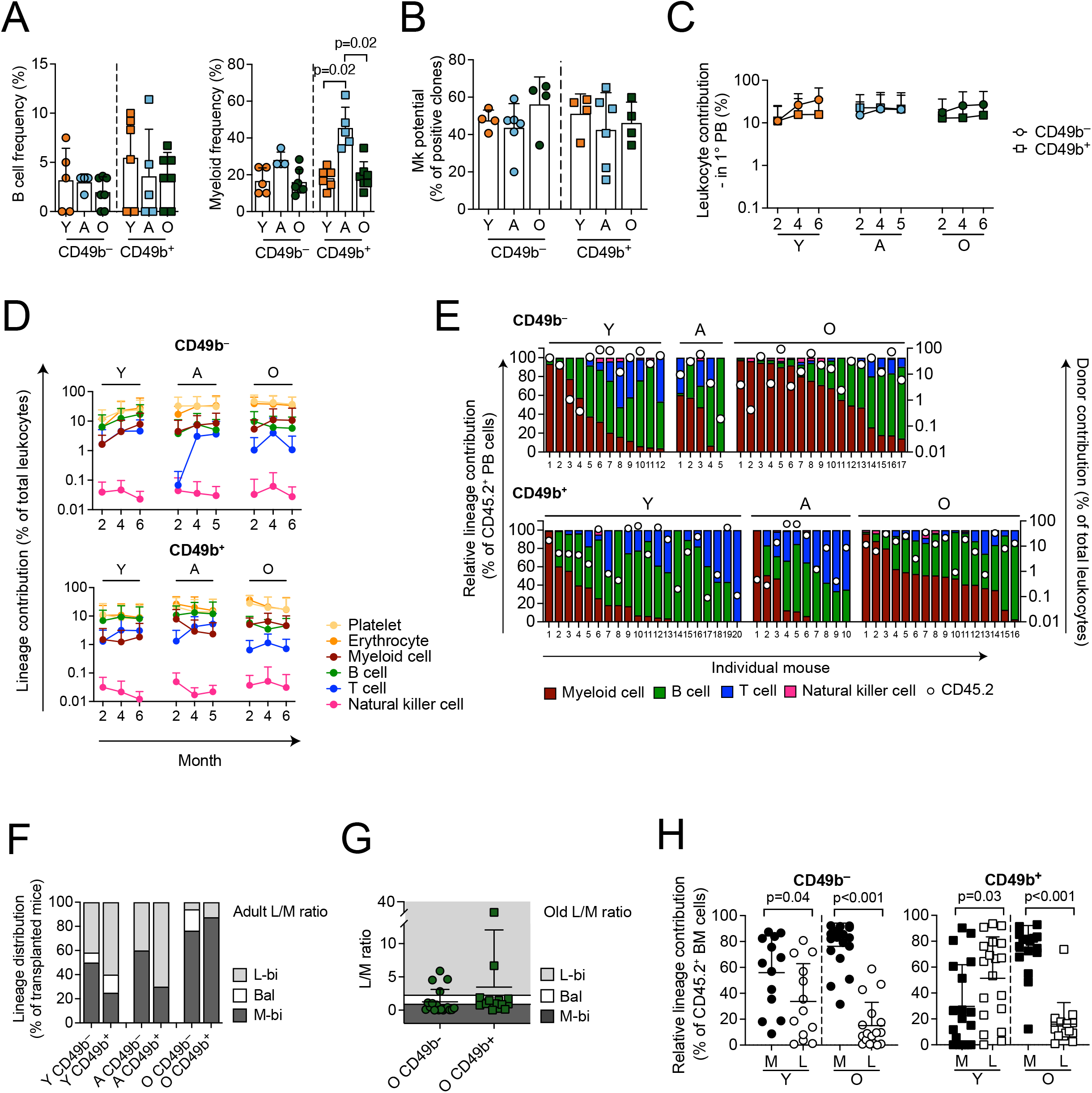
The lineage repopulation patterns of multipotent CD49b subsets change with age. **a**, Total frequency of clones with B cells and/or B and myeloid cells (left), and clones containing only myeloid cells (right) from single cell sorted CD49b^−^ and CD49b^+^ HSC subsets from young (n_CD49b_^−^ = 5, n_CD49b_^+^ = 6, 2 experiments), adult (n_CD49b_^−^ = 4, n_CD49b_^+^ = 5, 2 experiments), and old (n_CD49b_^−^ = 7, n_CD49b_^+^ = 7, 4 experiments) mice. **b**, Megakaryocyte differentiation potential of single cell sorted CD49b^−^ and CD49b^+^ HSC subsets from young (n_CD49b_^−^ = 4, n_CD49b_^+^ = 4, 2 experiments), adult (n_CD49b_^−^ = 6, n_CD49b_^+^ = 6, 5 experiments), and old (n_CD49b_^−^ = 4, n_CD49b_^+^ = 4, 4 experiments) mice. **c**, Total donor leukocyte contribution in the peripheral blood (PB) of mice transplanted with CD49b^−^ and CD49b^+^ HSC subsets from young, adult, and old mice (Y_CD49b_^−^ = 14, Y_CD49b_^+^ = 24, YA_CD49b_^−^= 5, YA_CD49b_^+^ = 10, O_CD49b_^−^ = 17, O_CD49b+_ = 16). **d**, Total donor-derived lineage contribution to platelets, erythrocytes, myeloid, B, T, and natural killer cells in the PB of mice transplanted with CD49b^−^ and CD49b^+^ HSC subsets from young, adult, and old mice (Y_CD49b_^−^ = 14, Y_CD49b_^+^ = 24, YA_CD49b_^−^ = 5, YA_CD49b_^+^ = 10, O_CD49b_^−^ = 17 (8 for P-E) and O_CD49b_^+^ = 16 (9 for P-E)). **e,** Relative lineage contribution to donor leukocytes (left axis) and total donor chimerism (right axis) in the PB of individual mice transplanted with CD49b^−^ and CD49b^+^ HSC subsets from young, adult, and old mice 5-6 months post-transplantation. **f,** Frequency of observed lineage distribution patterns of transplanted mice with CD49b^−^ and CD49b^+^ HSCs from young, adult, and old mice 5-6 months post-transplantation, from (e), calculated using lymphoid to myeloid (L/M) cell ratio in PB from adult unmanipulated mice as a reference (Y_CD49b_^−^ = 12, Y_CD49b_^+^ = 20, YA_CD49b_^−^ = 5, YA_CD49b_^+^ = 10, O_CD49b_^−^ = 17, O_CD49b_^+^ = 16). **g,** Calculated L/M cell ratio in PB, of mice transplanted with old CD49b^−^ and CD49b^+^ HSCs (O_CD49b_^−^ = 17, O_CD49b_^+^ = 16) at 5-6 months post-transplantation. The observed L/M ranges for lymphoid-biased (L-bi), balanced (Bal), and myeloid-biased (M-bi) based on unmanipulated old mice are indicated. See Extended Data Fig. 3d for L/M ratio in unmanipulated mice. **h,** Lineage contribution to myeloid (M) and lymphoid cells (L: B, T, and NK cells) in the bone marrow of mice transplanted with young and old CD49b^−^ and CD49b^+^ HSC subsets, 6 months post-transplantation (Y_CD49b_^−^ = 13, Y_CD49b_^+^ = 19, O_CD49b_^−^ = 17, O_CD49b_^+^ = 16). Mean ± s.d.is shown. The statistical analysis was performed with Kruskal-Wallis with Dunn’s multiple comparison test in (a-b), Mann-Whitney test in (c,h). Y, young; A, adult; O, old; PB, peripheral blood; BM, bone marrow; L-bi, lymphoid-biased; Bal, balanced; M-bi, myeloid-biased; L/M, lymphoid to myeloid. See also Extended Data Figs. 1-3.

To evaluate age-associated differences in multilineage differentiation ability of CD49b subsets *in vivo*, we performed competitive transplantation experiments using the *Gata-1* eGFP mouse strain to permit detection of platelets and erythrocytes, in addition to leukocytes^27^. A limiting dose of five cells from young or adult CD49b^−^ and CD49b^+^ subsets were transplanted into each recipient, while one hundred cells from old mice were transplanted to account for the reduced regenerative capacity of old HSCs^1, 2, 6^. As expected, the total leukocyte contribution of the old HSC subsets was comparable to their young and adult counterparts (Fig. 2c). Although both CD49b subsets from all age groups demonstrated multilineage repopulating ability, they exhibited subset-and age-dependent repopulation patterns (Fig. 2d and Extended Data Figs. 2 and 3a,b). Most notably, young CD49b^−^ cells, in contrast to adult and old CD49b^−^ cells, repopulated lymphoid cells, especially B cells, with high efficiency and at similar levels as the young CD49b^+^ cells (Fig. 2d). On the contrary, old CD49b^+^ cells repopulated myeloid cells more efficiently than young and adult CD49b^+^ cells, and at comparable level to the old CD49b^−^ subset (Fig. 2d). Therefore, to assess the relative contribution of lymphoid and myeloid cells within the donor leukocyte compartment, we analyzed the lineage distribution of individually transplanted mice (Fig. 2e). Using peripheral blood (PB) profiles from adult unmanipulated mice as reference (Extended Data Fig. 3c), the lineage distribution of transplanted mice was categorized based on the observed lymphoid to myeloid blood cell (L/M) ratio (Extended Data Fig. 3d)^22, 28^. In accordance with previous data^22^, the most frequent categorization in young and adult CD49b^−^ cells was the myeloid-biased (M-bi) pattern, whereas lymphoid-biased (L-bi) was most common in CD49b^+^ cells (Fig. 2f)^22^. Therefore, lineage distribution between young and adult mice was highly similar (Fig. 2d-f), consistent with observed steady-state blood profiles (Extended Data Fig. 3c,d). In stark contrast to young and adult age groups, both CD49b subsets in old mice were highly M-bi (Fig. 2f), consistent with the blood profiles of old unmanipulated mice (Extended Data Fig. 3c). Given the overall higher myeloid contribution in old mice, we used the L/M ratio from old unmanipulated mice as a reference to assess whether old CD49b^−^ and CD49b^+^ transplanted mice enriched for different lineage distribution patterns (Extended Data Fig. 3d). Classifying repopulation patterns based on the old L/M ratio revealed that old CD49b^+^ HSCs greatly enriched for cells with a balanced (Bal) cellular output, while M-bi was still the predominant classification in old CD49b^−^ HSCs (Fig. 2g and Extended Data Fig. 3e).

**Fig. 3.**
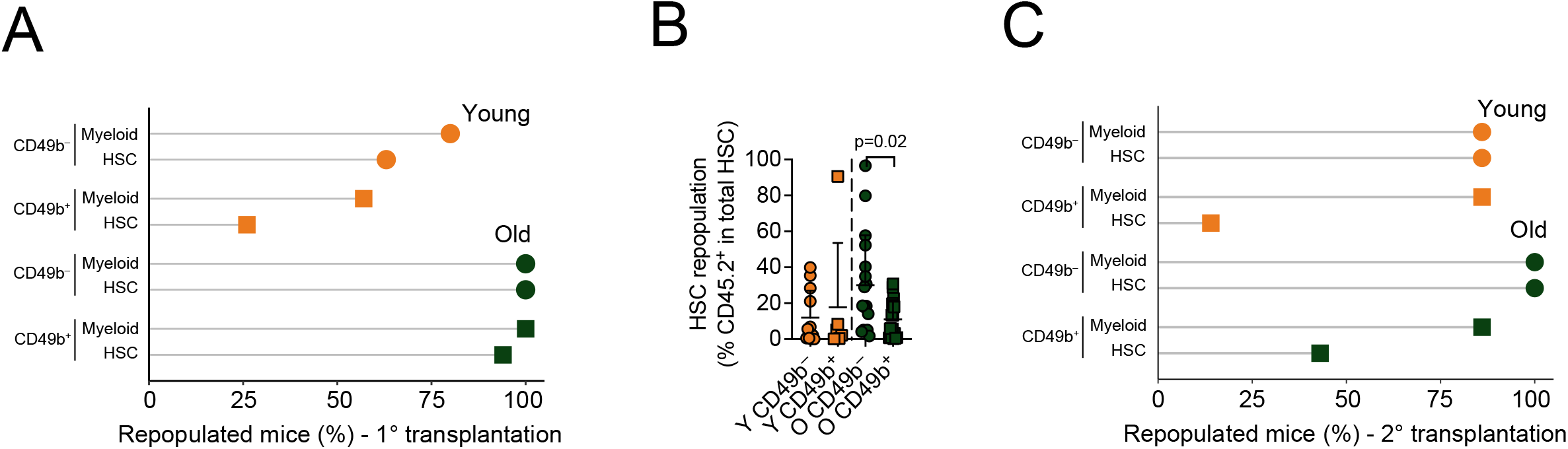
The CD49b^−^ HSC subset is the most durable subset regardless of age. **a**, Proportion of mice exhibiting myeloid repopulation in the peripheral blood and HSC (LSK CD48^−^Flt-3^−^ CD150^+^) repopulation in the bone marrow (BM), 5-6 months after primary transplantation of young or old CD49b^−^ and CD49b^+^ HSCs (Y_CD49b_^−^ = 16, Y_CD49b_^+^ = 23, O_CD49b_^−^ = 17, O_CD49b_^+^ = 16). **b,** Frequency of HSC repopulation in reconstituted mice from (a). Mean ± s.d. is shown. The statistical analysis was performed with Mann-Whitney test. Y, young; O, old. **c,** Proportion of mice exhibiting myeloid repopulation in the peripheral blood and HSC (LSK CD48^−^Flt-3^−^CD150^+^) repopulation in the BM, 6 months after primary transplantation of young or old CD49b^−^ and CD49b^+^ HSCs (Y_CD49b_^−^ = 7, Y_CD49b_^+^ = 7, O_CD49b_^−^ = 2, O_CD49b_^+^ = 7). See also Extended Data Fig. 4.

In accordance with blood repopulation patterns, the young CD49b^−^ population mainly generated myeloid cells and the young CD49b^+^ subset potently generated lymphocytes in the BM compartment (Fig. 2h). In contrast, both old CD49b^−^ and CD49b^+^ subsets had higher relative myeloid repopulation in the BM (Fig. 2h). Furthermore, nearly all mice transplanted with old CD49b^−^ and CD49b^+^ HSCs repopulated the granulocyte-monocyte progenitors (GMPs) and megakaryocyte progenitors (MkPs), whereas only a few mice reconstituted the common lymphoid progenitors (CLPs) and lymphoid-primed multipotent progenitors (LMPPs) (Extended Data Fig. 4a-d).

**Fig. 4.**
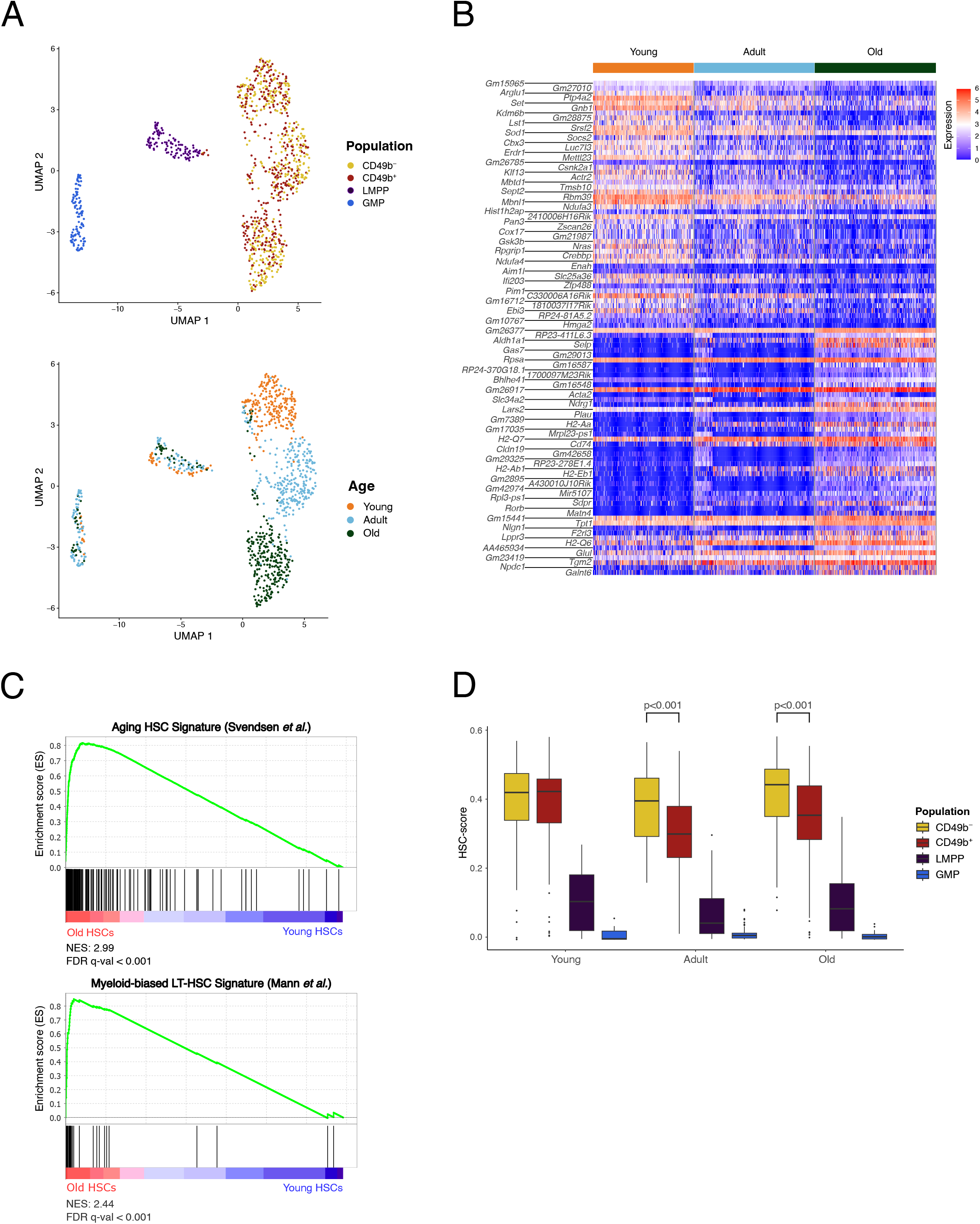
HSCs undergo considerable gene expression changes during aging. **a**, UMAP visualization of single-cell RNA-seq data from stem-and progenitor cells, from young (Y), adult (A), and old (O) mice (Y_CD49b_^−^ = 115, Y_CD49b_^+^ = 118, Y_LMPP_ = 34, Y_GMP_ = 11, A_CD49b_^−^ = 133, A_CD49b_^+^ = 144, A_LMPP_ = 55, A_GMP_ = 73, O_CD49b_^−^ = 145, O_CD49b_^+^ = 135, O_LMPP_ = 32, O_GMP_ = 26). Cells are colored by population (top) or age group (bottom). **b,** Heatmap of normalized expression for differentially expressed genes between young and old HSCs. The top 50 differentially expressed genes up in young and old, with p_adj_ < 0.01 and log_2_FC > 1, are shown. **c,** Gene set enrichment analysis for old compared to young HSCs and the indicated custom gene sets from Svendsen *et al*.^33^ and Mann *et al*.^36^ NES, normalized enrichment score. **d,** Calculated HSC-score for stem-and progenitor cells from young, adult, and old mice. The statistical analysis was performed with Mann-Whitney test. See also Extended Data Fig. 5.

Overall, our findings demonstrate that both CD49b^−^ and CD49b^+^ cells increase their myeloid repopulation with age, and that the CD49b cell surface marker distinguishes different functional HSC subsets throughout aging.

### The CD49b^−^ HSC subset is the most durable subset regardless of age

Different cell populations within the HSC compartment were shown to differ in self-renewal ability^11, 29, 30^. Hematopoietic stem cell subsets fractionated with CD49b in adult mice were previously shown to differ in the durability of their self-renewal potential^22^. Therefore, to investigate the changes of extensive self-renewal ability of CD49b^−^ and CD49b^+^ subsets in aging, we assessed the number of mice that actively reconstituted myeloid cells in the PB 5-6 months post-transplantation, as a measure of ongoing HSC activity. Long-term (LT) myeloid repopulation was more frequently found in mice transplanted with young CD49b^−^ compared to young CD49b^+^ cells (Fig. 3a). Consistent with the differences observed in LT myeloid repopulating activity, the number of transplanted mice which reconstituted phenotypic HSCs was higher in the young CD49b^−^ compared to the young CD49b^+^ group, but with no difference in their overall repopulation level (Fig. 3a,b). Furthermore, young CD49b^−^ and CD49b^+^ HSCs were able to regenerate both types of phenotypic HSC subsets (Extended Data Fig. 4e). In contrast, all reconstituted mice from the old age group had LT myeloid repopulation with no difference between subsets, compatible with the high frequency of mice that also repopulated phenotypic HSCs, likely reflecting the higher number of transplanted cells (Fig. 3a). However, although both old CD49b populations were able to regenerate phenotypic HSC subsets, the old CD49b^−^ subset reconstituted HSCs more efficiently than the old CD49b^+^ subset (Fig. 3b and Extended Data Fig. 4f). These results suggest that CD49b subsets differ in their durable self-renewal ability in young and old mice. Therefore, to examine the self-renewal potential of young and old CD49b^−^ and CD49b^+^ subsets more conclusively, transplanted mice which exhibited phenotypic HSC repopulation were secondary transplanted. Concordant with previous studies, the repopulation patterns were generally preserved through serial transplantation (Extended Data Fig. 4g,h)^22, 28^. Although more than 80% of secondary transplanted mice from young and old CD49b populations repopulated myeloid cells long term, CD49b^−^ subsets from young and old mice more frequently reconstituted phenotypic HSCs compared to their CD49b^+^ counterparts (Fig. 3c). Our findings are compatible with CD49b^−^ HSCs harboring the highest self-renewal potential in both young and old mice.

### HSCs undergo considerable gene expression changes during aging

To investigate the molecular mechanisms underlying functional differences of HSC subsets in aging, we performed single cell RNA-sequencing (scRNA-seq)^31, 32^ on stem-and progenitor cells from young and old mice (CD49b^−^, CD49b^+^, LMPP and GMP; supplementary Table 1), and combined this with our previously published data from adult mice^22^.

Using dimensionality reduction, we observed that HSCs, LMPPs and GMPs formed distinct clusters (Fig. 4a), with expected expression of genes used for phenotypic definition of the populations, including *Slamf1*, *Flt3, Cd48*, *Fcgr2b* and *Fcgr3* (Extended Data Fig. 5a). While the CD49b subsets were not distinguishable, the HSCs segregated into age-specific clusters, in stark contrast to the progenitors (Fig. 4a). We subsequently investigated differentially expressed genes between young and old HSCs (CD49b^−^ and CD49b^+^ combined; p_adj_<0.01, log_2_FC>1) and found 173 genes upregulated and 455 genes downregulated with age (Fig. 4b and supplementary Table 1). Several reported aging-related genes were observed, including *Selp* (P-selectin) and *Aldh1a1* (aldehyde dehydrogenase 1a1; Extended Data Fig. 5a)^18, 33–35^. Furthermore, gene set enrichment analysis (GSEA) demonstrated that genes associated with HSC aging and myeloid lineage bias had increased expression in old HSCs (Fig. 4c)^33, 36^.

**Fig. 5.**
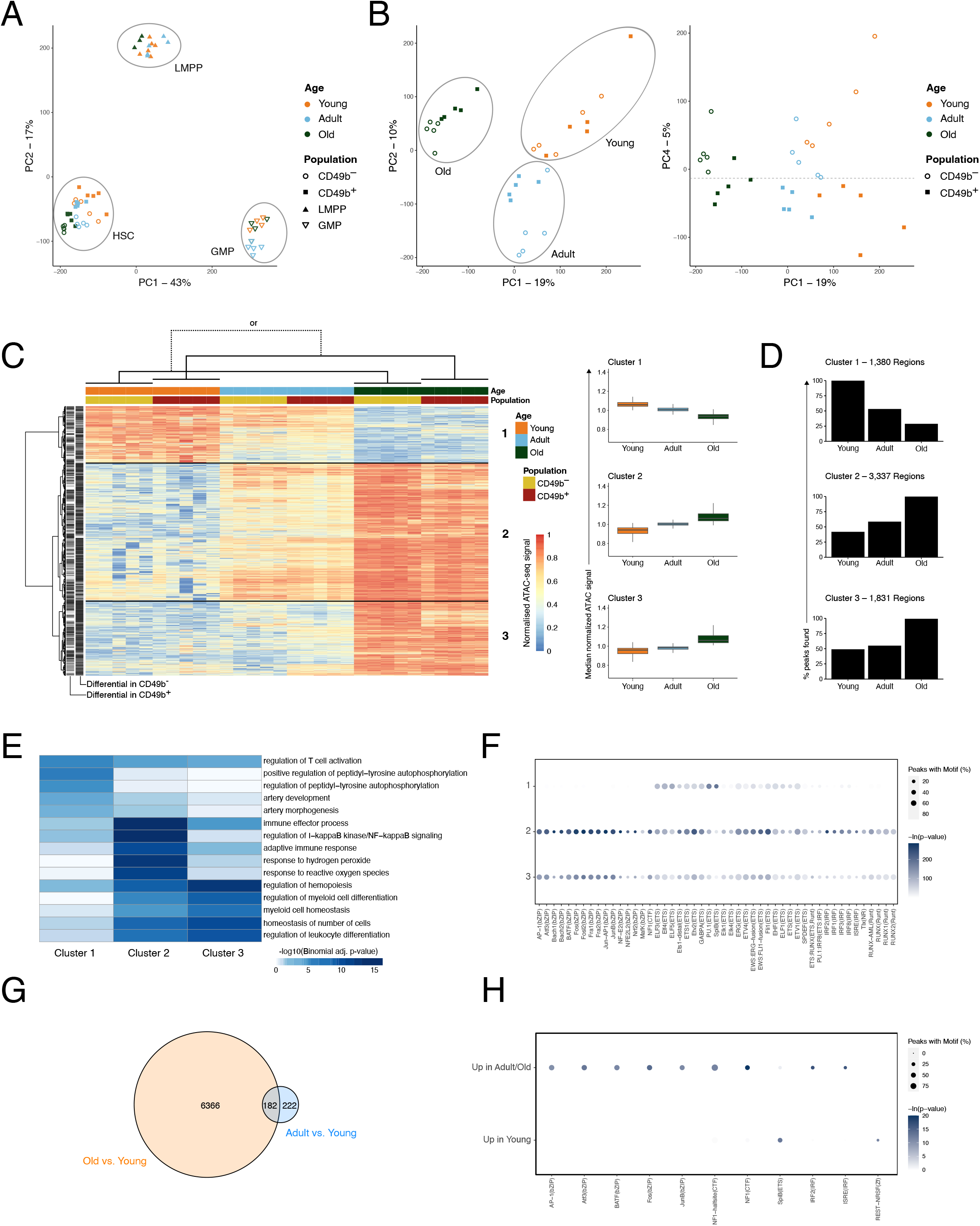
Aging is associated with a progressive increase of chromatin accessibility in HSCs. **a**, Principal component analysis of ATAC-seq data from stem-and progenitor cells, from young (Y), adult (A), and old (O) mice (Y_CD49b_^−^ = 5, Y_CD49b_^+^ = 5, Y_LMPP_ = 6, Y_GMP_ = 4, A_CD49b_^−^ = 5, A_CD49b_^+^ = 5, A_LMPP_ = 5, A_GMP_ = 5, O _CD49b–_ = 5, O_CD49b_^+^ = 5, O_LMPP_ = 3, O_GMP_ = 4). **b,** Principal component analysis of ATAC-seq data from CD49b^−^ and CD49b^+^ HSC subsets, from young (Y), adult (A), and old (O) mice. (Y _CD49b–_ = 5, Y_CD49b_^+^ = 5, A_CD49b_^−^ = 5, A_CD49b_^+^ = 5, O _CD49b–_ = 5, O_CD49b_^+^ = 5). **c,** Heatmap (left) of row normalized chromatin accessibility for regions with differential accessibility (p_adj_<0.0001) between young and old CD49b^−^ and/or between young and old CD49b^+^ cells. Regions are divided into three clusters based on hierarchical clustering. Median normalized chromatin accessibility of clusters 1-3 are shown (right). **d,** Percentage of regions constituting open chromatin in clusters 1-3. **e,** Top 5 GO biological processes significantly enriched in clusters 1-3. **f,** Transcription factors with enriched binding motifs (-ln(p-value)>50) in clusters 1-3. **g,** Venn diagram of regions with differential accessibility (p_adj_<0.0001) in old compared to young HSCs (Old vs. Young) or in adult compared to young HSCs (Adult vs. Young). **h,** Transcription factors with enriched binding motifs (-ln(p-value)>10) in regions with increased or decreased accessibility in both adult and old compared to young HSCs. See also Extended Data Figs. 5-6.

In line with the transcriptional similarity between CD49b^−^ and CD49b^+^ cells (Fig. 4a), no individual genes were found to be significantly different between them in young and old age groups (p_adj_<0.05). However, given the distinct functional differences between CD49b subsets, we therefore applied the hscScore method, which utilizes validated HSC gene expression data sets^37^ to assess whether CD49b subsets can be molecularly distinguished by leveraging potentially small but concordant differences in gene expression. Notably, we observed a significantly lower HSC-score in adult and old CD49b^+^ compared to the corresponding CD49b^−^ cells, in agreement with the reduced self-renewal potential of CD49b^+^ subsets (Figs. 3 and 4d)^22^. In contrast, both CD49b^−^ and CD49b^+^ cells from young mice had a similar score, consistent with their comparable HSC repopulating efficiency (Figs. 3b and 4d).

Collectively, these results demonstrate that aging is associated with distinct gene expression changes in HSCs, which interestingly is not observed in progenitors. Moreover, global transcriptome analysis cannot resolve functionally distinct HSC subsets, even at the single cell level^22^.

### Aging is associated with a progressive increase of chromatin accessibility in HSCs

Given the high transcriptional similarity between CD49b^−^ and CD49b^+^ cells, we next used Assay for Transposase-Accessible Chromatin sequencing (ATAC-seq)^38^ to examine age-related epigenetic differences in stem-and progenitor cells from young and old mice (supplementary Table 2 and Extended Data Fig. 5b,c). Our previously published data from adult mice were included for comparison^22^. Genes such as *Slamf1*, *Flt3*, and *Fcgr2b*, showed expected population specific chromatin accessibility profiles (Extended Data Fig. 5d). Principal component analysis (PCA) of the ATAC-seq data revealed distinct clusters of LMPPs, GMPs, and HSCs (CD49b^−^ and CD49b^+^; Fig. 5a), concordant with scRNA-seq data. Although HSC subsets overall created one cluster (Fig. 5a), discrete subclusters based on age group and CD49b surface expression were discernable in principal components (PC) 1-3 (Fig. 5b, left and Extended Data Fig. 6a) and PC4 (Fig. 5b, right), respectively.

**Fig. 6.**
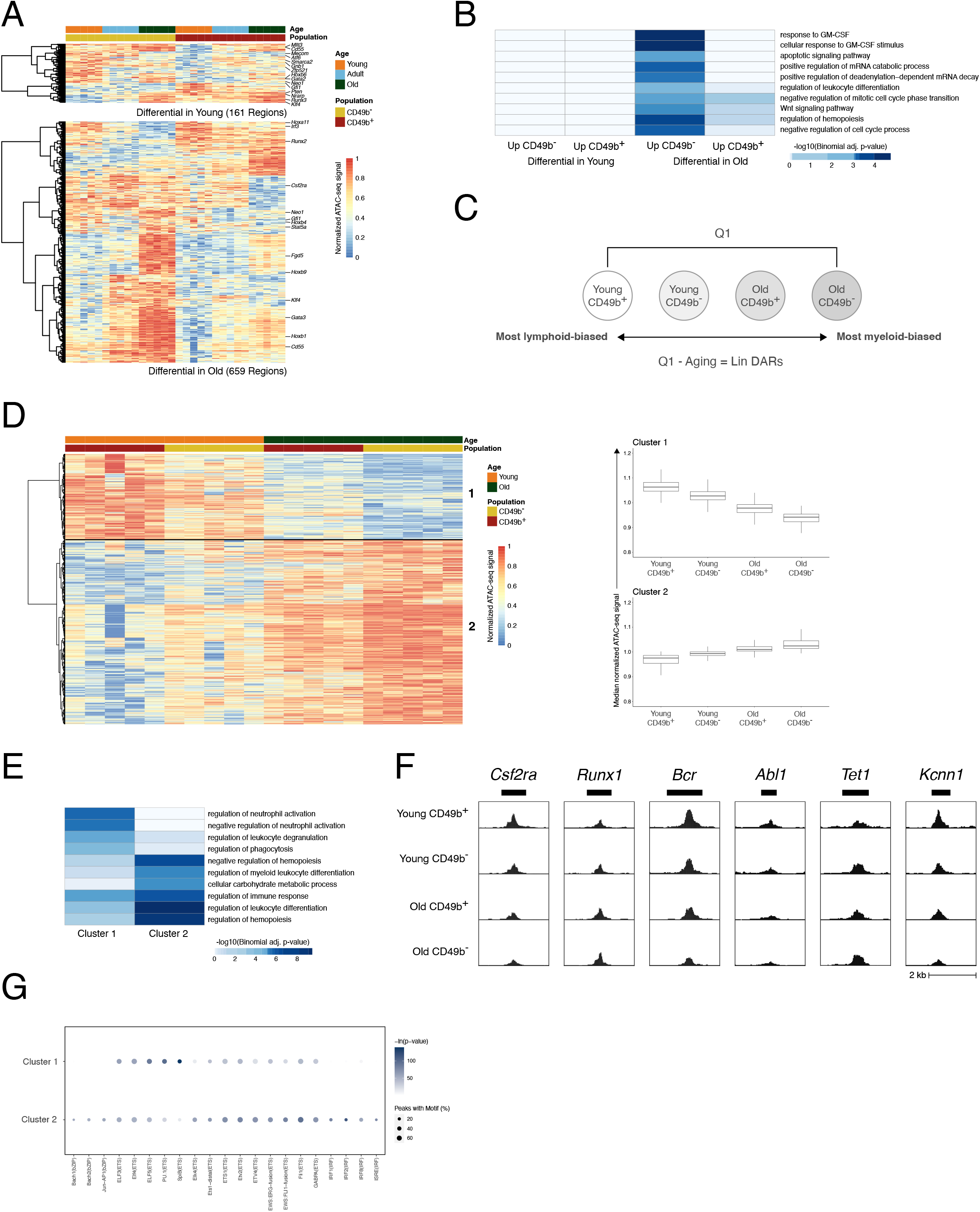
Aging and lineage bias are regulated by the same transcription factor families. **a**, Heatmap of row normalized chromatin accessibility for regions with differential accessibility (p_adj_<0.05) between young CD49b^−^ and CD49b^+^ HSCs (top), or between old CD49b^−^ and CD49b^+^HSCs (bottom). **b,** Top 10 GO biological processes significantly enriched in regions with differential accessibility between CD49b subpopulations in young or old mice. **c,** Schematic illustration of the analysis strategy to identify chromatin accessibility changes associated with lineage bias differences. Lin DARs, lineage bias associated differentially accessible regions. **d,** Heatmap (left) of row normalized chromatin accessibility for Lin DARs. Regions are divided into two clusters based on hierarchical clustering. Boxplots (right) show the median normalized chromatin accessibility in clusters 1 and 2. **e,** Top 5 GO biological processes significantly enriched in clusters 1 and 2. **f,** UCSC browser tracks of median ATAC-seq signal for selected Lin DARs. Gene names above the tracks indicate the closest gene to the displayed region. **g,** Transcription factors with enriched binding motifs (-ln(p-value)>50) in clusters 1 and 2. See also Extended Data Fig. 6.

As the PCA indicated substantial age-related differences in HSCs, we next interrogated the chromatin accessibility changes between young and old CD49b subsets. We identified 5,501 and 3,849 significantly differentially accessible regions (DARs; p_adj_<0.0001) between young and old CD49b^−^, and between young and old CD49b^+^ subsets, respectively (Fig. 5c and supplementary Table 2). The majority of DARs constituted age-associated gain in accessibility, including regions of common age-related genes including *Clu* (Clusterin), *Aldh1a1*, and *Cdc42* (Cell division cycle 42; Extended Data Fig. 6b)^33, 35, 39^. Overall, the pattern of age-dependent chromatin accessibility changes was highly similar in both CD49b subsets and was generally initiated already in adult HSCs (Fig. 5c). These findings suggest common aging mechanisms in both subsets. Strikingly, in old HSCs, regions with gained chromatin accessibility were largely acquired *de novo*, whereas regions with decreased accessibility frequently lost accessibility completely (Fig. 5d). Interestingly, the chromatin accessibility changes in HSCs are reversible, as age-associated changes were not propagated to downstream progenitors to any large extent (Extended Fig. 6c).

Gene ontology (GO) analysis of DARs with high accessibility in young HSCs (Fig. 5c, cluster 1) was associated with T cell activation and cell signaling (Fig. 5e). Conversely, DARs with high accessibility in old HSCs (Fig. 5c, clusters 2-3) enriched for processes connected to reactive oxygen species (ROS) and NF-κB signaling responses (Fig. 5e, cluster 2), myeloid cell differentiation processes, hematopoiesis, and cell number regulation (Fig. 5e, cluster 3). These results are consistent with the functional decline in lymphopoiesis and increased myelopoiesis with age, as well as increased NF-κB signaling and ROS accumulation in aging^35, 40–43^.

We next performed motif enrichment analysis to assess putative transcription factor (TF) binding in aging HSCs. Notably, DARs with increased accessibility in young HSCs (Fig. 5c, cluster 1) predominantly enriched for transcription factor binding sites (TFBS) of the ETS family TFs, of which SPI1 (PU.1) and SPIB were the most significantly enriched in young compared to adult and old HSC subsets (Fig. 5f). In contrast, DARs with increased accessibility in adult and old HSCs (Fig. 5c, clusters 2-3) were enriched for ETS-, as well as bZIP-, IRF-, and RUNT- family TFBS (Fig. 5f). In agreement with the motif enrichment analysis, *Spi1* (*PU.1*) and *Spib* gene expression was reduced in aging, whereas expression of *Junb*, *Irf1*, *and Runx1* increased with age (Extended Data Fig. 7a).

**Fig. 7.**
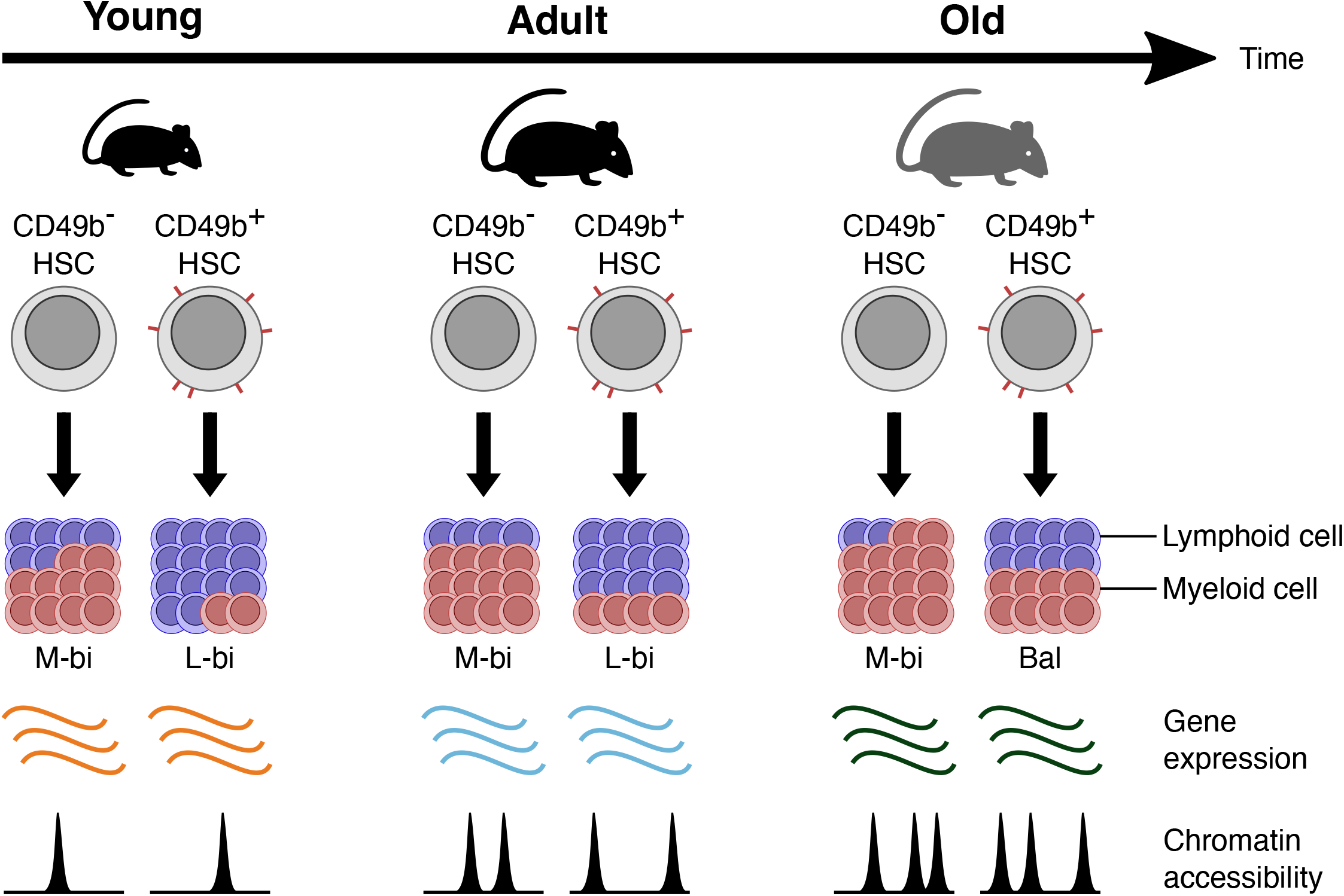
Aging is associated with functional and molecular changes in distinct hematopoietic stem cell subsets. Schematic overview of the age-related functional and molecular changes in CD49b^−^ and CD49b^+^ HSCs in young, adult, and old mice. Aging is associated with gradually increasing myeloid cellular output in both CD49b^−^ and CD49b^+^ HSCs. Gene expression profiling detects age-related molecular changes in total HSCs. Chromatin accessibility analysis reveals age-dependent and CD49b subset specific differences. M-bi, myeloid-biased; L-bi, lymphoid-biased; Bal, balanced.

We observed a gradual increase of open chromatin with age, with adult HSC subsets exhibiting intermediate chromatin accessibility (Fig. 5c) that was already initiated in the transition from young to adult HSCs. Therefore, to further investigate the early aging-associated chromatin regions we assessed the overlap of DARs between young and adult HSCs, and between young and old HSCs (Fig. 5g). We identified 182 age-related DARs that were already significantly changed from young to adult stages (p_adj_<0.0001). Motif enrichment analysis of the shared regions showed enrichment of TFBS from bZIP-, CTF-, and IRF-families in DARs with gained accessibility in aging. Conversely, SPIB TFBS were enriched in DARs with reduced accessibility with age (Fig. 5h).

Collectively, our findings demonstrate that aging is primarily associated with an HSC-specific and progressive gain of chromatin accessibility in both CD49b^−^ and CD49b^+^ subsets that is already initiated in the transition from young to adult stage. Furthermore, our data suggest that aging-related chromatin remodeling is linked to age-dependent changes in TF binding.

### Aging and lineage bias are regulated by the same transcription factor families

Although the age-related chromatin changes in CD49b^−^ and CD49b^+^ subsets were highly similar (Fig. 5c), the subsets clustered apart within all age groups in the PCA (Fig. 5b). By investigating the chromatin accessibility differences between CD49b^−^ and CD49b^+^ subsets in young and old mice, we identified 161 and 659 DARs (p_adj_<0.05), respectively (Fig. 6a and supplementary Table 2). The high similarity in chromatin accessibility between CD49b^−^ and CD49b^+^ HSCs in young mice corresponds to their comparable cell cycling and proliferation characteristics, as well as HSC repopulating activity (Extended Data Fig. 1e,g and Fig. 3a,b). In further agreement, GO analysis of DARs between young CD49b subsets did not yield any significantly enriched terms (Fig. 6b). In contrast, the old CD49b^−^ group enriched for processes including cell cycle and leukocyte regulation, as well as GM-CSF and WNT pathways (Fig. 6b). These findings are consistent with old CD49b^−^ cells being more quiescent and myeloid-biased compared to old CD49b^+^ cells (Fig. 2g and Extended Data Fig. 1e,g).

In transplantation experiments, we observed considerable functional differences in blood lineage contribution, with an overall increase in myeloid differentiation in both CD49b subsets with age. To identify epigenetic changes associated with lineage bias differences, we first categorized HSC subsets based on their differentiation characteristics (Fig. 6c). Differential analysis was subsequently performed between the most lymphoid-biased (young CD49b^+^) and myeloid-biased (old CD49b^−^) population to identify DARs associated with lineage bias. However, given the substantial chromatin changes in aging HSCs (Fig. 5c,d), we filtered out aging-related DARs (Fig. 5c and supplementary Table 2) to identify DARs uniquely associated with lineage bias (Lin DARs, Fig. 6c,d and supplementary Table 2). The Lin DARs separated into a smaller cluster with high accessibility regions in lymphoid-biased young CD49b^+^ HSCs and a larger cluster signified by higher accessibility in myeloid-biased old CD49b^−^ HSCs (Fig. 6d). GO analysis revealed enrichment of several blood cell associated processes, including regulation and differentiation of leukocytes, myeloid cells, and neutrophils, confirming that the identified chromatin regions are important for lineage differentiation (Fig. 6e). In further agreement, the Lin DARs included regions associated with the myeloid growth factor receptor, *Csf2ra* (colony stimulating factor 2 receptor, alpha) and hematopoietic regulator, *Runx1* (runt related transcription factor 1) (Fig. 6f and Extended Data Fig. 7b)^44, 45^. Additionally, we identified other candidate genes within the Lin DARs with a potential role in regulating HSC lineage differentiation (supplementary Table 2). For example, we found *Bcr* (BCR activator of RhoGEF and GTPase) and *Abl1* (c-abl oncogene 1, non-receptor tyrosine kinase) genes, which are necessary for normal neutrophil and lymphoid cell function respectively (Fig. 6f; Extended Data Fig. 7b). Intriguingly, chromosomal translocation of t(9;22)(q34;q11) results in the fusion oncogene BCR-ABL1, which is characteristic of the HSC-derived chronic myelogenous leukemia, but also present in acute lymphoblastic leukemia^46–48^. Furthermore, we identified *Tet1* (tet methylcytosine dioxygenase 1), which is suggested as a negative regulator of HSC self-renewal and B cell differentiation, and with a tumor suppressor role in B cell lymphoma^49, 50^. Unexpectedly, we also detected *Kcnn1* (potassium intermediate/small conductance calcium-activated channel, subfamily N, member 1), which is mainly expressed in the brain but also associated with the bone tumor Ewing sarcoma^51, 52^. However, the role of *Kcnn1* in hematopoiesis is not well-characterized.

To identify candidate TFs influencing HSC lineage bias and differentiation, we performed motif enrichment analysis of cluster 1 and 2 Lin DARs (Fig. 6g). Intriguingly, the enriched TFBS belonged to the same TF families as the aging related TFs, including ETS-, bZIP-, IRF-, and RUNT-families. Notably, SPI1 (PU.1) and SPIB TFBS were highly enriched in young CD49b^+^ HSCs, whereas bZIP-, IRF-, and RUNT-TFBS were most significantly enriched in old CD49b^−^ HSCs.

Altogether, our results suggest that CD49b subsets in young mice are both functionally and epigenetically similar but become more distinctly different with age. Furthermore, our findings indicate that aging and lineage bias may largely be governed by the same TFs.

## Discussion

As the global population live longer, health implications due to aging have become a public health concern. Age is a risk factor for neurodegenerative and chronic diseases, as well as cancer^4^. Physiologic aging of the hematopoietic system is associated with perturbed immunity and impaired homeostasis, leading to the onset of anemia and increased risk of blood malignancies^1–3^. Notably, there is a predominance of myeloid malignancies with increasing age, whereas the incidence of lymphoid malignancies is higher in children and young people^2, 53, 54^. It has been suggested that the age-dependent impairment of HSC function may play a role in aging of the hematopoietic system^16^. Therefore, insights into age-related HSC behavior are critical to understand and overcome physiological consequences of an aging hematopoietic system.

The overall predominance of myeloid cells in aging was first proposed to be due to the functional decline of HSCs to produce lymphoid cells. With the identification of functionally diverse HSC subsets with differential blood lineage preferences, a different model was suggested, in which a change of HSC clonal composition underlies the myeloid dominance^1, 2^. However, many aging studies investigating molecular mechanisms of HSC aging have assessed the composite HSC compartment containing multiple diverse HSC subtypes, but age-associated features of highly enriched HSC subsets with distinct behavior are incompletely elucidated. Furthermore, most aging studies using mouse models compare adult (2-4 months) and old (1.5-2 years) age groups. However, HSCs switch from a fetal to an adult HSC phenotype already around 1 month after birth. This juvenile period involves active tissue growth and development in young mice^26^, which could contribute to the higher likelihood of lymphoid malignancies in children^54, 55^, highlighting the importance of encompassing the pre-adult stage of development when investigating age-associated changes in HSC behavior.

In this study, we have isolated CD49b^−^ and CD49b^+^ subsets^22^ from the phenotypic HSC compartment and investigated their functional and molecular changes throughout aging using young, adult, and old mice (Fig. 7). Although we did not find any significant changes in the proportion of CD49b^−^ and CD49b^+^ cells with age, functional assessment revealed distinct biological differences. Remarkably, there was a general shift towards the myeloid lineage with age, which was progressive and occurred in both myeloid-biased and lymphoid-biased enriched HSC subsets. Consequently, lineage distribution was altered, resulting in a more pronounced dominance of M-bi repopulation patterns in CD49b^−^ HSCs and a switch from mainly a L-bi to a Bal pattern in CD49b^+^ cells with age. Our data suggest that the CD49b^−^ subset enriches for myeloid-biased HSCs in all age groups, while the CD49b^+^ subset enriches for lymphoid-biased HSCs in young and adult mice, but marks lineage-balanced HSCs in old mice. These results suggest that age-related myeloid dominance in the hematopoietic system is due to intrinsic alterations in lineage differentiation properties of all HSCs, although we cannot exclude the possibility that there might also be changes in the HSC composition.

It has been demonstrated that lineage bias is a heritable trait,^28^ indicating that epigenetic regulatory mechanisms may underlie HSC heterogeneity^56^. We have previously shown that functionally different HSCs exhibit similar gene expression patterns but distinct epigenetic profiles. Our scRNA-seq analysis captured clear age-related gene expression changes but did not detect any global differences between HSC subsets, consistent with the importance of gene regulatory mechanisms governing HSC heterogeneity^22^. Indeed, we observed both age-and subset-related epigenetic differences by ATAC-seq analysis. We found that chromatin accessibility progressively increased in both HSC subsets, which correlated with the loss of SPIB and SPI1 (PU.1) TFBS, consistent with previous reports^18, 21, 57^. Interestingly, we found that reduced SPIB TFBS were already detected in adult HSCs, indicating that age-related remodeling of chromatin regions is initiated early in the transition from the young to adult stage. Our data agree with the hypothesis that some gene expression changes in aging may originate from the juvenile period as part of the growth restricting process that occurs into adulthood^58^. Our results also implicate that aging-induced chromatin changes could be used to prospectively identify aging epigenetic signatures. Remarkably, the age-related molecular changes were primarily observed in HSCs and not propagated to progenitors, suggesting that aging mechanisms preferentially target HSCs. Further functional investigations are necessary to establish whether interference of chromatin remodeling in these regions could be exploited to reverse or remedy aging alterations in HSCs.

Although our ATAC-seq data showed that age-dependent changes were more substantial than changes between CD49b^−^ and CD49b^+^ HSCs of the same age group, subset-specific epigenetic differences were still detected, which became more distinct in old mice. Strikingly, the same TF families were enriched in both age-related and subset-specific chromatin regions. Indeed, the ETS family members SPI1 (PU.1) and SPIB, which were identified to be downregulated in aging, have also been demonstrated to be important for myeloid and lymphoid development and differentiation^59, 60^. This finding highlights the possibility that aging and lineage differentiation processes may involve the same TFs. Our ATAC-seq analysis also identified several candidate genes with possible roles in regulating HSC lineage bias. Among these, *Bcr*, *Abl1*, and *Tet1* have been shown to be necessary for proper myeloid and lymphoid lineage differentiation, whereas *Kcnn1* has not been studied in the hematopoietic system yet. Deficiencies or aberrant activation of *Bcr*, *Abl1*, and *Tet1* cause hematopoietic malignancies, highlighting the importance of elucidating the regulatory mechanisms of normal blood lineage differentiation^46–52^. Further functional studies are needed to clarify the involvement of *Bcr*, *Abl1*, *Tet1*, and *Kcnn1* in HSC lineage bias regulation.

Our data suggest that HSC subsets with the same immunophenotype gradually change their functional and epigenetic attributes during aging, starting already in the transition from the young to the adult stage. The intrinsic changes, including alterations in lineage preference, quiescence, and proliferation states, are correlated with significant remodeling of the chromatin landscape. Our findings are therefore compatible with aging models suggesting cell-aautonomous HSC changes^2, 17^. Collectively, we have demonstrated that CD49b resolves HSC subsets with age-dependent cellular and epigenetic changes (Fig. 7). Our studies provide important insights into the contribution of distinct HSC subsets and the consequences of their functional alterations to the changing hematopoietic compartment during aging. Clarifying the role of HSC subsets in aging is critical towards understanding and providing therapeutic prospects to overcome age-associated dysfunction of the hematopoietic system, including malignancies.

## Methods

### Animals

Female and male C57BL/6J mice used in the experiments were housed and maintained at the Karolinska University Hospital Preclinical Laboratory, Sweden. Young mice between 3 and 4 weeks old, adult mice between 7 and 18 weeks old, and old mice between 77 and 117 weeks old (1.5-2 years) were used. C57BL/6J or Gata-1 eGFP mice,^27^ backcrossed >8 generations to C57BL/6J (CD45.2), were used as donors and B6.SJL-*Ptprc^a^Pepc^b^/*BoyJ mice (CD45.1) were used as primary and secondary recipients in transplantation experiments. All experiments were approved by the regional ethical committee, Linköping ethical committee in Sweden (ethical numbers: 882 and 02250-2022).

### Transplantation experiments

In primary transplantation experiments, donor cells (CD45.2^+^) were sorted and intravenously injected with 200,000 CD45.1^+^ support BM cells into irradiated adult CD45.1 mice. Five sorted HSCs from young and adult donor mice and 100 HSCs from old donor mice were transplanted. The full irradiation dose was given in two doses with at least 4 hours in between each dose by a ^137^*Cesium source.* A total of 10 Gy was used in experiments with young and old donors and total 12 Gy in experiments with adult donors.

In secondary transplantation experiments, 10·10^6^ unfractionated BM cells from phenotypic HSC (LSK CD48^−^CD150^+^) reconstituted primary recipient mice were intravenously injected into 1-5 lethally irradiated recipients. Recipient mice were monitored and PB analyses were done regularly up to 5-6 months post-transplantation for both primary and secondary transplantation experiments.

### Preparation of hematopoietic cells

Bone marrow (BM) single cell suspensions were prepared by crushing femurs, tibiae, and iliac crests isolated from the mice into Phosphate-Buffered Saline (PBS, Gibco) supplemented with 5% Fetal Bovine Serum (Gibco) and 2mM Ethylenediaminetetraacetic acid (EDTA, Merck).

Unfractionated BM cells were counted on the XP-300-Hematology Analyzer (Sysmex Corporation) and then Fc-blocked either with purified CD16/32 (BD Biosciences), or stained with CD16/32 antibody conjugated to a fluorophore. Following Fc-block, the cells were stained with antibodies against cell surface marker antigens. See Supplementary Table 3 for antibodies used in flow cytometry analyses and Supplementary Table 4 for phenotypic definitions of hematopoietic populations.

To detect hematopoietic stem cells (HSCs), unfractionated BM cells were enriched with CD117 MicroBeads (Miltenyi Biotec), and subsequently selected using immunomagnetic separation before subsequent staining with antibodies against cell surface markers.

Peripheral blood (PB) was sampled from transplanted mice from the tail vein. Blood was collected in lithium heparin coated microvette tubes (Sarstedt). The platelet fraction was separated from whole blood samples by centrifugation, and leukocytes were subsequently isolated with Dextran sedimentation. Isolated platelets, erythrocytes, and leukocytes were stained with antibodies against cell surface antigens as described previously.^22, 61^

### Flow Cytometry analysis of hematopoietic cells

Flow cytometry analyses were done on FACSymphony™ A5 and LSR Fortessa™ following cell preparation. For cell sorting experiments, FACSAria™ Fusion cell sorters (BD Biosciences) were used. The mean cell sorting purity was 93%±6%, calculated from 28 experiments. Fluorescence minus one (FMO) controls were included in every experiment. For single cell *in vitro* experiments, single cell sorting into individual wells of 96-well or 72-well plates were confirmed by sorting 488-nm fluorescent beads (ThermoFisher Scientific). Analyses following data acquisition were done using FlowJo software version 10 (BD Biosciences).

### Calculation of reconstitution and lineage bias

Donor reconstitution was calculated based on the frequency of CD45.2^+^ events in total leukocytes. Transplanted mice with ≥0.1% total donor contribution in leukocytes (CD45.2^+^) and/or platelets in the peripheral blood (PB), represented by ≥10 events in the donor gate of the recipient mice, were scored as positively repopulated at month 2 post-transplantation.

Blood lineage repopulation was calculated based on the frequency of CD45.2^+^ events in the leukocyte lineages or of Gata-1 eGFP^+^ events in platelet and erythrocytes. Transplanted mice were determined to be positive for a specific blood lineage when the repopulation was ≥0.01% and represented by ≥10 events in the donor gate.

Relative donor reconstitution levels were calculated based on the frequency of B, T, NK, and myeloid cells within the CD45.2^+^ cells at month 5-6 post-transplantation.

The blood lineage distribution of the leukocyte fraction was calculated based on the ratio of lymphoid (L) to myeloid (M) cells (L/M) in the PB of adult or old unmanipulated mice. Lymphoid cells included B, T, and NK cells. The calculated L/M ratios from unmanipulated mice (Extended Data Fig. 3d) were used to categorize the lineage distribution in the PB of transplanted mice at month 5-6 post-transplantation.

### *In vitro* assays

Myeloid (CD11b^+^Gr-1^+^ and/or F4/80^+^CD11b^+^) and B cell (CD19^+^B220^+^) differentiation potential was assessed using the OP9 co-culture assay by sorting single cells onto OP9 stroma, which was analyzed after 3 weeks of culture. Megakaryocyte potential was assessed by single cell sorting into 72-well plates (ThermoFisher Scientific) and evaluated after 11 days by scoring the presence of megakaryocytes using an inverted microscope. Cell division kinetics were carried out by tracking cell divisions of single sorted cells into 60-well plates (ThermoFisher Scientific) for 3 days post-sort. See Supplementary Table 5 for culture conditions.

### Cell cycle and proliferation assays

The cell cycle state of HSCs was analyzed using Ki-67 staining, following the manufacturer’s protocol from the BD Cytofix/Cytoperm Kit (BD Biosciences). The cell proliferative state of HSCs was assessed using 5-Bromo-2’-deoxyuridine (BrdU) incorporation, where one dose of BrdU was given by intraperitoneal injection (50mg/g bodyweight, BD Biosciences), followed by oral administration via drinking water (800mg/mL, Merck) for 3 days post-injection. The BrdU experimental process and visualization were performed according to the BrdU Flow Kit protocol (BD Biosciences).

### Single cell RNA- and ATAC-sequencing

Single cells were deposited into 384-well plates for SmartSeq2 single cell RNA-seq^32^ as previously described^22, 31^. Libraries were sequenced on HiSeq3000 (Illumina) using dual indexing and single 50 base-pair reads. See Supplementary Table 1 for sequenced RNA-seq samples.

Reads were demultiplexed, aligned to the mm10 reference genome using Tophat2 and deduplicated using samtools. For further analysis, R and the Seurat package (v.4.3.0) were used. Reads mapping to *CT010467.1* were excluded as these reads are assumed to largely originate from rRNA contamination. Cells were filtered to have an RNA count between 50,000 and 750,000 reads, <10% mitochondrial reads, and <10% ERCC spike-in contribution. Lowly expressed genes with ≤400 reads across all included cells were filtered out. Data were normalized using Seurat’s SCTransform function, regressing out the percentage of mitochondrial reads. PCA was run on the 3,000 most variable genes and the top 10 principal components were used as input to create UMAP plots. Differential expression analysis was done with Seurat’s FindMarkers function using a logistic regression framework (LR test) and the percentage of mitochondrial reads as latent variable. Gene set enrichment analysis (GSEA) was run using the GSEA desktop application and custom gene sets obtained from relevant literature. HSC-scores were calculated using the hscScore tool^37^ using the provided Jupyter notebook and trained model, as suggested by the developers. Statistical analysis of HSC-scores was done in R using the Mann-Whitney test.

Bulk ATAC-seq was performed with 500 sorted cells from C57BL/6J mice of young and old age groups using a modified Omni-ATAC protocol, as previously described^22^. Samples were then amplified and paired-end sequencing (2 x 41 cycles) was performed on NextSeq 500 (Illumina). See Supplementary Table 2 for sequenced ATAC-seq samples.

Sequence reads were aligned to the mm10 reference genome and subsequently filtered using the nf-core ATAC-seq pipeline (v1.0.0). The nf-core pipeline was also used for peak calling with MACS2. Samples were quality checked and poor-quality samples (Fraction of Reads in Peaks, FRiP, less than 10%) were excluded. Read coverage was normalized to 10^6^ mapped reads in peaks for all samples and for each population, median normalized read coverage was calculated and visualized using the UCSC genome browser. Read positions were adjusted by +4bp and −5bp for the positive and negative strand, respectively. HOMER^62^ was used to annotate peaks to the closest gene, as well as to quantify reads in consensus peaks. Peaks with more than 5 fragments per kilobase million (FPKM) in at least a third of the samples were considered true/found in a population, while peaks not found in any of the populations were filtered and excluded. R was subsequently used to log transform and quantile normalize read counts for visualization. Following peak calling and filtering, differentially accessible regions were determined using the DESeq2 package in R. Differential regions not found in either compared population were excluded from further analysis. Adult ATAC-seq data obtained from Somuncular *et al.*^22^ were included during the bioinformatic analysis with young and old HSC ATAC-seq samples. To match the sample size and sample quality from the young and old age groups, only 5 adult samples with the highest FRiP scores for each cell population were included in the analysis. GREAT^63^ (http://great.stanford.edu/public/html/<u>, v4.0.4</u>) was used for gene ontology analysis using online default settings to determine significant enrichment. Motif enrichment analysis was done using HOMER.

### Statistical analysis

GraphPad Prism v.9.4.1 for Mac OS or R was used for all statistical analyses. Non-parametric tests were performed with the Mann-Whitney or the Kruskal-Wallis with Dunn’s multiple comparison. Mean ± SD and *p*-values are indicated in all figures.

## Data availability

ATAC-seq and scRNA-seq data have been deposited in the European Nucleotide Archive (ENA) with accession number PRJEB55627.

## Author contributions

Contribution: E.S., T.-Y.S., Ö.D., A.-S.J. and S.L. performed experiments and analyzed data; C.G., E.T., and A.F. assisted in experiments; J.H., and T.-Y.S. performed bioinformatics analysis; A.-S.J, R.M., and S.L. supervised the work; S.L. designed the project; S.L. wrote the paper together with E.S., J.H., T.-Y.S., A.-S.J and R.M.; all authors reviewed the manuscript before submission.

## Competing interests

The authors declare no competing financial interests.

## Acknowledgements

The authors thank Claus Nerlov (University of Oxford) for providing the Gata-1 eGFP mouse strain; Joakim Dillner (Karolinska Institutet, KI) for sequencing support; Preclinical Laboratory, Karolinska University Hospital and KI MedH Flow Cytometry Core Facility for their services; Hui Gao, KI Centre for Bioinformatics and Biostatistics (CBB) for helpful advice; KI Single cell core Facility @ Flemingsberg campus (SICOF) for their single cell sequencing services. The computations and data storage were enabled by resources provided by the National Academic Infrastructure for Supercomputing in Sweden (NAISS) and the Swedish National Infrastructure for Computing (SNIC) at Uppmax partially funded by the Swedish Research Council (2022- 06725 and 2018-05973). S.L was supported by a Wallenberg Academy Fellow award (2016.0131) and by the Swedish Childhood Cancer Fund (TJ2017-0074, PR2017-0047). J.H and T-Y.S were supported by KI Doctoral Education grants. This work was supported by grants from the European Hematology Association, the Swedish Cancer Society (CAN2017/583, 20 1062 PjF), the Swedish Research Council (2016-02331), the Strategic research area (SFO) in Stem cell and Regenerative Medicine, Åke Olsson foundation, Åke Wiberg foundation and King Gustav V Jubilee Fund.

---

**Extended Data Fig. 1, related to Figs. 1-2.**
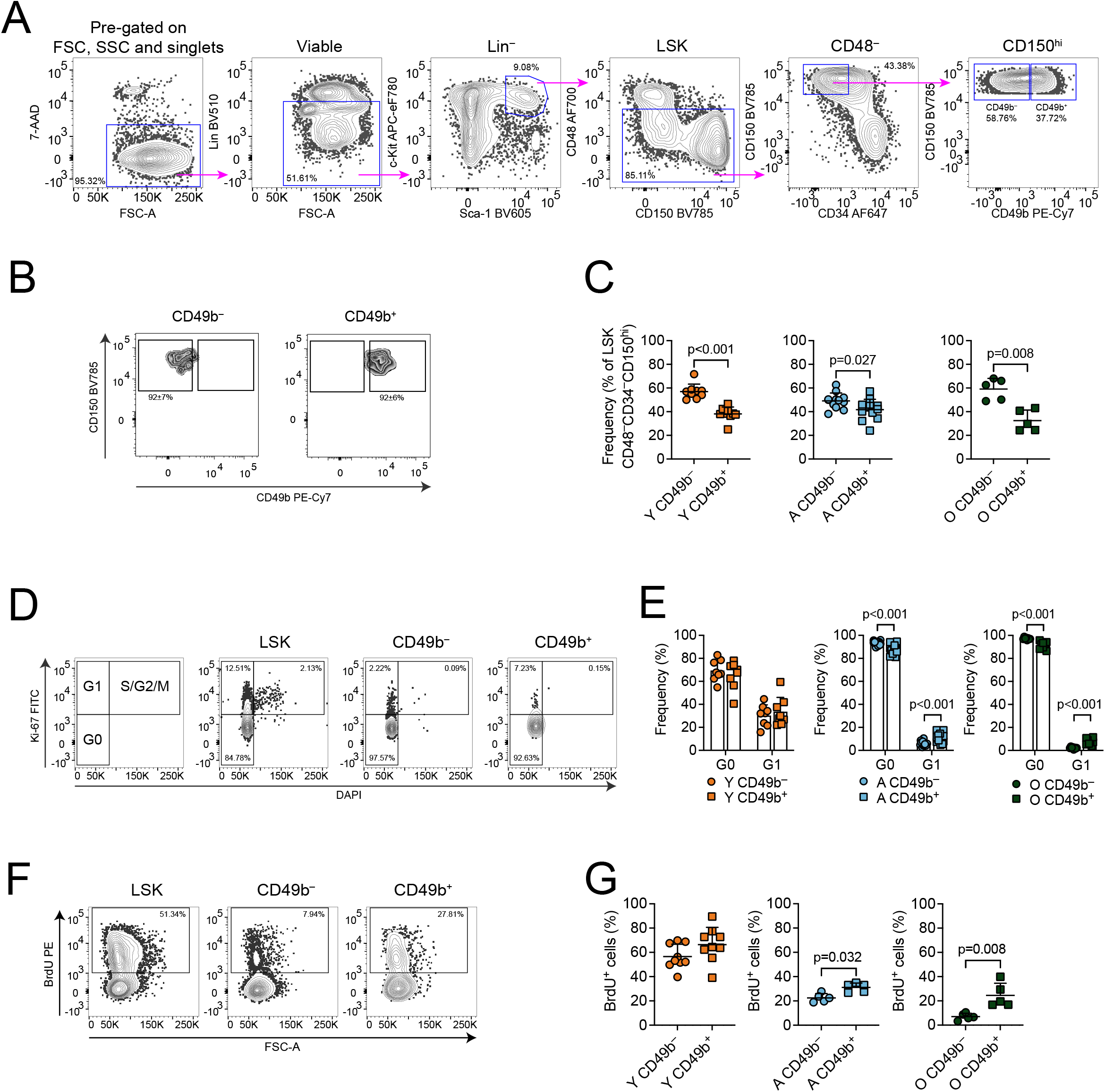
Fluorescence-activated cell sorting and functional evaluation of CD49b^−^ and CD49b^+^ HSC subsets in aging. **a**, Representative FACS profile and gating strategy of the phenotypic HSC compartment (Lin^−^Sca-1^+^c-Kit^+^ (LSK) CD48^−^CD34^−^CD150^hi^) and further separation with CD49b. Frequency of parent gates are shown. **b,** Sort purity analysis of CD49b^−^ and CD49b^+^ populations pre-gated on Lin^−^Sca-1^+^c-Kit^+^CD48^−^CD34^−^CD150^hi^. Sort purity is represented as mean ± s.d. for each subset, from 28 experiments. **c,** Frequency of CD49b^−^ and CD49b^+^ HSC subsets within the LSK CD48^−^ CD34^−^CD150^hi^ population in young (n = 9, 2 experiments), adult (n = 12, 6 experiments), and old (n = 5, 5 experiments) mice. **d,** Representative FACS profile of Ki-67 cell cycle analysis. **e,** Frequency of CD49b^−^ and CD49b^+^ HSCs in G0 and G1 of young (n = 8, 2 experiments), adult (n = 15, 6 experiments), and old (n = 8, 5 experiments) mice. **f,** Representative FACS profile of BrdU cell proliferation analysis. **g,** Frequency of BrdU^+^ CD49b^−^ and CD49b^+^ HSCs of young (n = 9, 2 experiments), adult (n = 5, 1 experiment), and old (n = 5, 2 experiments) mice. Mean ± s.d. is shown. The statistical analysis was performed with Mann-Whitney test. Y, young; A, adult; O, old.

**Extended Data Fig. 2, related to Fig. 2.**
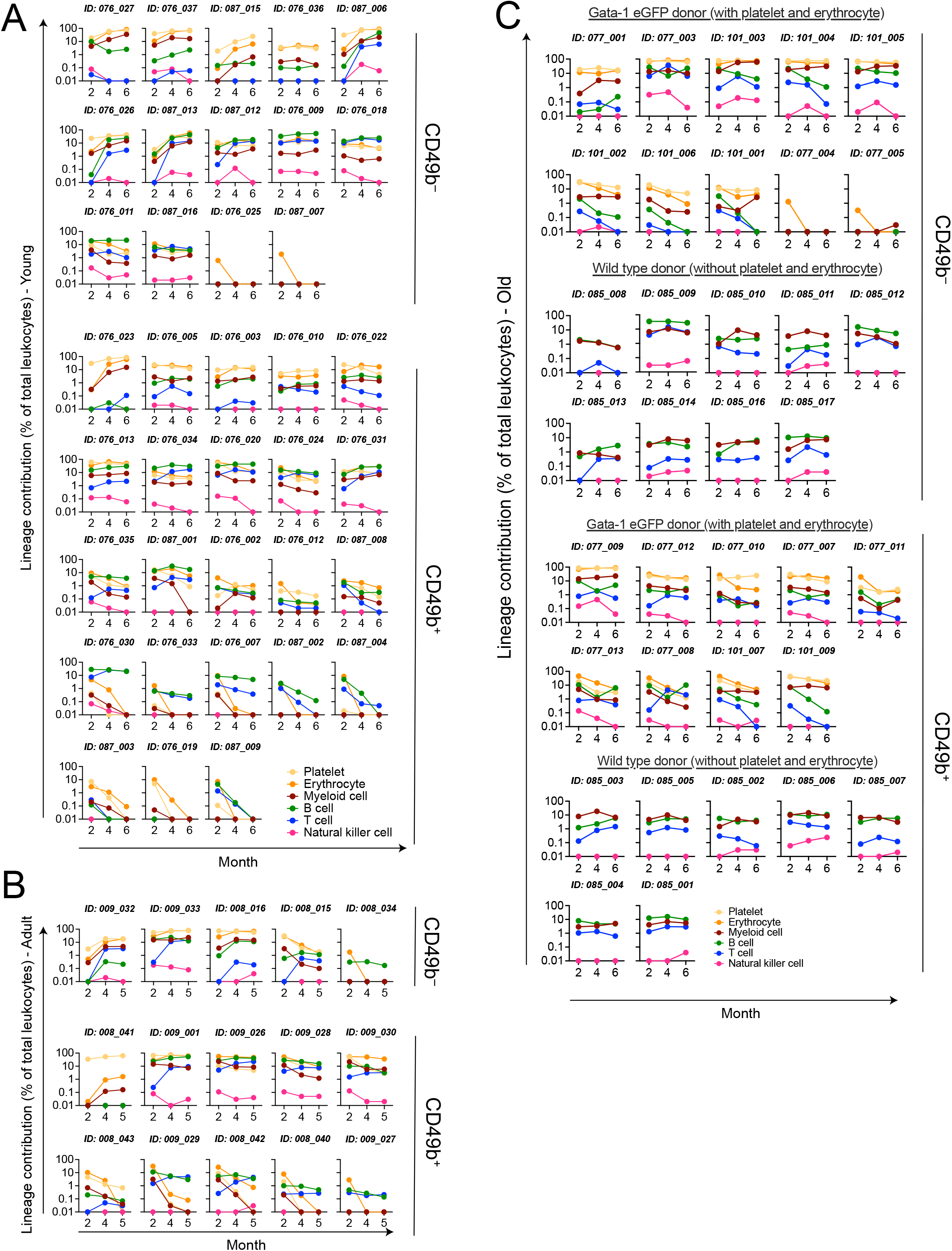
Repopulation profiles of primary transplanted mice from young, adult, and old HSC subsets. **a,** Total donor-derived blood lineage contribution in the peripheral blood of individual mice transplanted with CD49b^−^ and CD49b^+^ HSC subsets from young mice in primary transplantation (Y_CD49b_^−^ = 14 and Y_CD49b_^+^ = 23, 2 experiments). **b,** Total donor-derived blood lineage contribution in peripheral blood of individual mice transplanted with CD49b^−^ and CD49b^+^ HSC subsets from adult mice in primary transplantation (YA_CD49b_^−^ = 5 and YA_CD49b_^+^ = 10, 2 experiments). **c,** Total donor-derived blood lineage contribution in peripheral blood of individual mice transplanted with CD49b^−^ and CD49b^+^ HSC subsets from old mice in primary transplantation (O_CD49b_^−^ = 19 and O_CD49b_^+^ = 16, 3 experiments). Transplanted mice are grouped according to transplantation with wildtype C57BL/6J or Gata-1 eGFP donor mice.

**Extended Data Fig. 3, related to Fig. 2.**
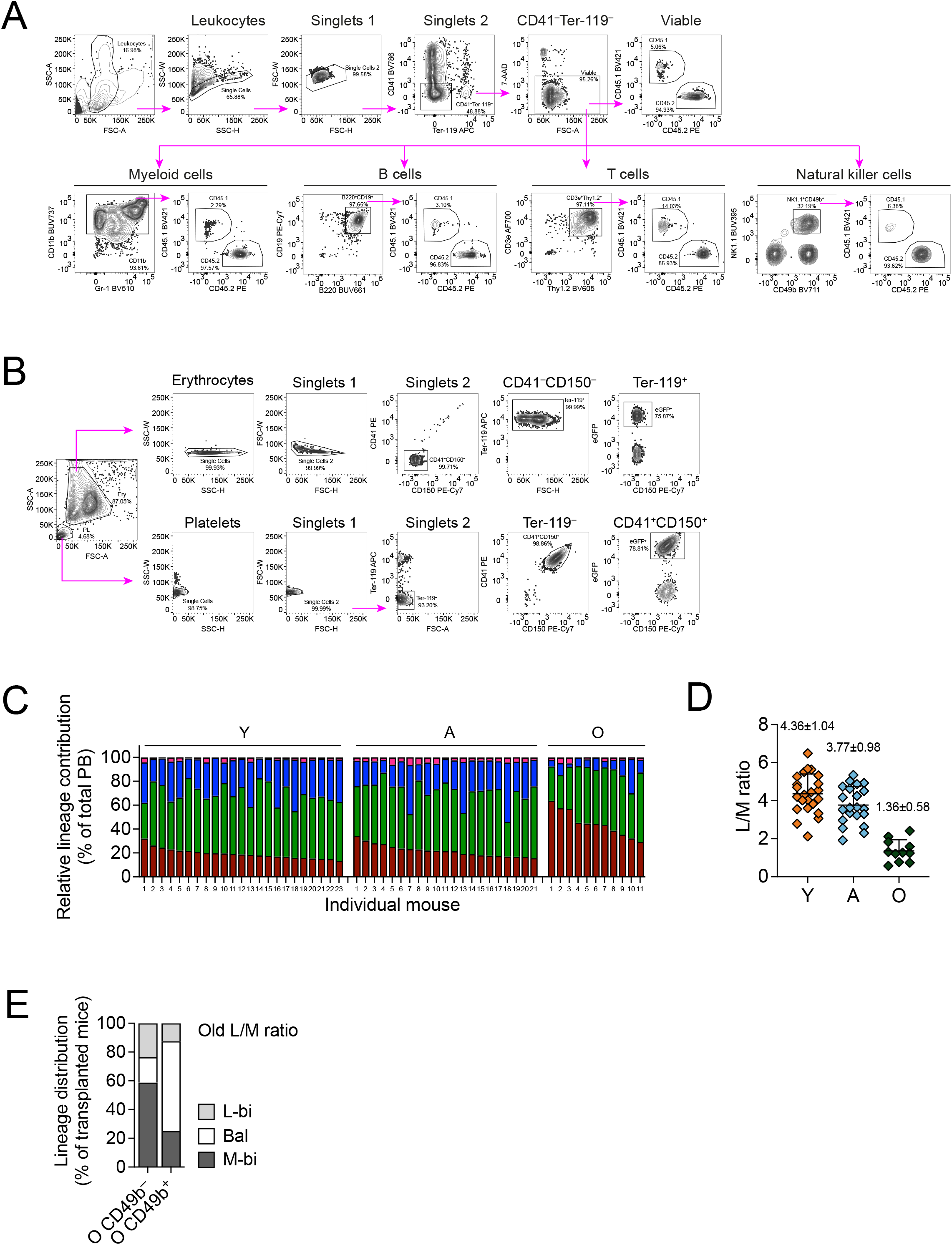
Flow cytometry analysis of mature blood lineages and blood lineage profiles in unmanipulated mice. **a**, Representative FACS profile and gating strategy of donor-derived myeloid cells, B cells, T cells, and natural killer cells in the peripheral blood. Frequency of parent gates are shown. **b,** Representative FACS profile and gating strategy of donor-derived platelets and erythrocytes in the peripheral blood. Frequency of parent gates are shown. **c,** Relative contribution of myeloid, B, T, and natural killer cells in peripheral blood leukocytes of individual unmanipulated young (n = 23, 3 experiments), adult (n = 21, 5 experiments), and old (n = 11, 1 experiment) mice. **d,** Calculated lymphoid (L: B, T, and NK cells) to myeloid (M) cell ratio in the peripheral blood based on unmanipulated young, adult, and old mice from (c). Mean ± s.d. is shown. **e,** Frequency of observed lineage distribution patterns of transplanted mice with CD49b^−^ and CD49b^+^ HSCs from old mice, 5-6 months post-transplantation, calculated using lymphoid/myeloid (L/M) cell ratio from old unmanipulated mice as a reference (O_CD49b_^−^ = 17, O_CD49b_^+^ = 16). PB, peripheral blood; Y, young; A, adult; O, old; L/M, lymphoid to myeloid; L-bi, lymphoid-biased; Bal, balanced; M-bi, myeloid-biased.

**Extended Data Fig. 4, related to Fig. 3.**
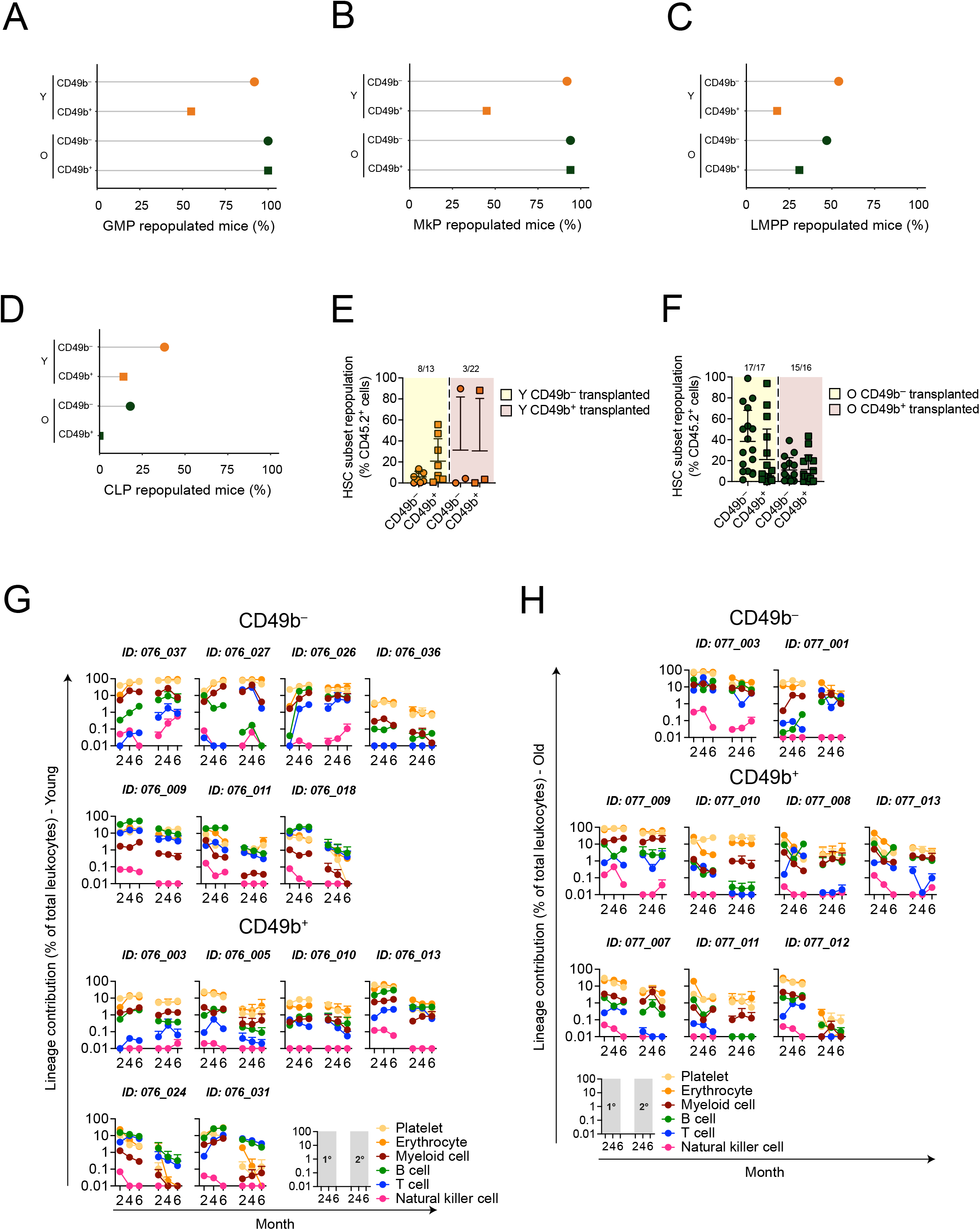
Repopulation of stem-and progenitor cell populations and secondary transplanted mice from young and old HSC subsets. **a-d**, Proportion of mice with GMP, MkP, LMPP, and CLP repopulation in the bone marrow (BM), 6 months after primary transplantation of young or old CD49b^−^ and CD49b^+^ HSCs. **e,** Frequency of phenotypic CD49b^−^ and CD49b^+^ HSC repopulation in the BM of reconstituted mice transplanted with young CD49b^−^ and CD49b^+^ HSC subsets in primary transplantation 6 months post-transplantation. **f,** Frequency of phenotypic CD49b^−^ and CD49b^+^ HSC repopulation in the BM of reconstituted mice transplanted with old CD49b^−^ and CD49b^+^ HSC subsets, in primary transplantation, 6 months post-transplantation. **g,** Total donor-derived blood lineage contribution in the peripheral blood (PB) of individual mice transplanted with CD49b^−^ and CD49b^+^ HSC subsets from young mice in primary and secondary transplantations (Y_CD49b_^−^ = 7 and Y_CD49b_^+^ = 6, 1 experiment, each primary donor mouse transplanted into 2-3 secondary recipients). **h,** Total donor-derived blood lineage contribution in PB of individual mice transplanted with CD49b^−^ and CD49b^+^ HSC subsets from old mice in primary and secondary transplantations (O_CD49b_^−^ = 2 and O_CD49b_^+^ = 7, 1 experiment, each primary donor mouse transplanted into 1-5 secondary recipients). In (e,f) the number of reconstituted primary donor mice out of all mice analyzed is indicated above the graphs. The statistical analysis was performed with the Mann-Whitney test. In (a-f): Y_CD49b_^−^ = 13, Y_CD49b_^+^ = 22, O_CD49b_^−^ = 17, O_CD49b_^+^ = 16. Mean ± s.d. is shown in (e-h). Y, young; O, old; GMP, granulocyte-monocyte progenitor; MkP, megakaryocyte progenitor; LMPP, lymphoid-primed multipotent progenitor; CLP, common lymphoid progenitor.

**Extended Data Fig. 5, related to Figs. 4-5.**
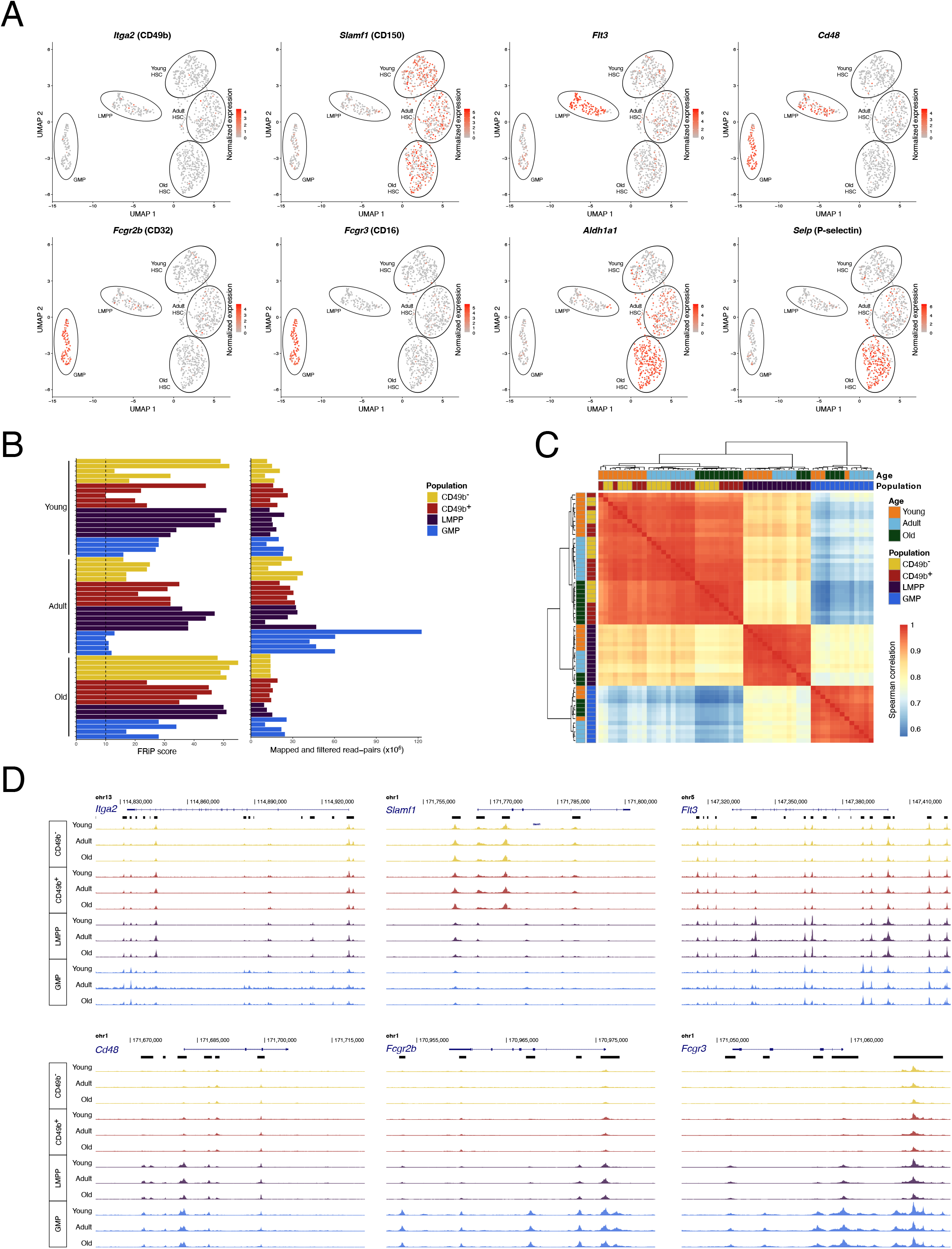
Gene expression and chromatin accessibility of selected genes and ATAC-seq sample characteristics. **a**, UMAP visualization of gene expression of selected, characteristic cell surface markers and aging-associated genes by scRNA-seq. The color intensity represents normalized expression level. **b,** Distribution of fraction of reads in peaks (FRiP) score and number of mapped and filtered read-pairs among ATAC-seq samples. **c,** Spearman correlation between ATAC-seq samples. Correlation is calculated on log_10_ transformed and quantile normalized data. **d,** UCSC browser tracks of median ATAC-seq signal for selected, characteristic cell surface markers for the studied cell populations.

**Extended Data Fig. 6, related to Fig. 5.**
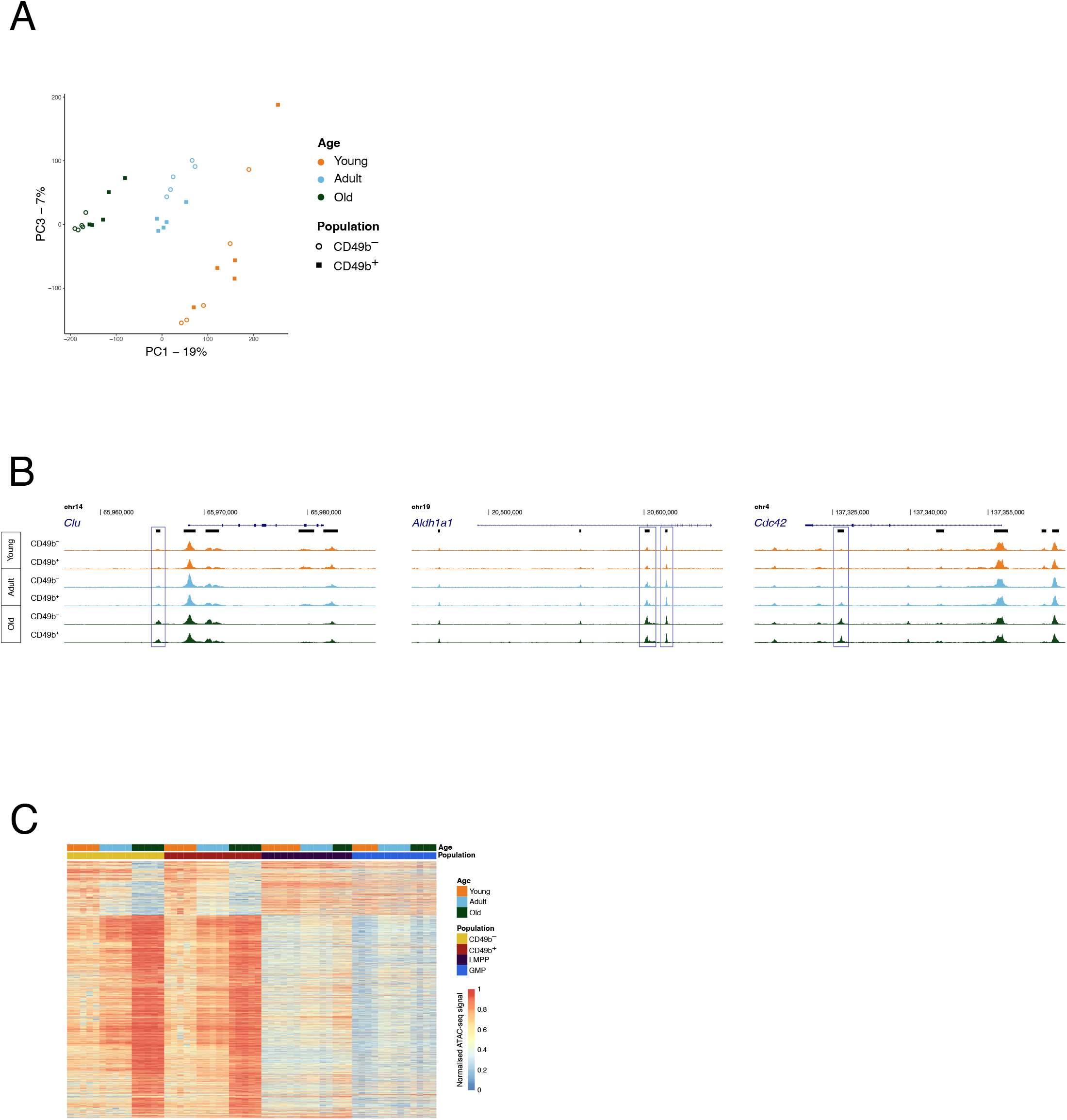
Gene expression and chromatin accessibility of selected gene regions associated with aging. **a**, Principal component analysis of ATAC-seq data from CD49b^−^ and CD49b^+^ HSC subsets, from young (Y), adult (A), and old (O) mice (Y_CD49b_^−^ = 5, Y_CD49b_^+^ = 5, A_CD49b_^−^ = 5, A_CD49b_^+^ = 5, O _CD49b–_ = 5, O_CD49b_^+^ = 5). Principal components 1 and 3 are shown. **b**, UCSC browser tracks of median ATAC-seq signal for selected regions. Peaks with significantly changed accessibility between young and old HSCs are indicated with blue boxes. **c,** Heatmap of row normalized chromatin accessibility for regions with differential accessibility (p_adj_<0.0001) between young and old CD49b^−^ and/or between young and old CD49b^+^ cells. LMPPs and GMPs are included in the heatmap.

**Extended Data Fig. 7, related to Fig. 6.**
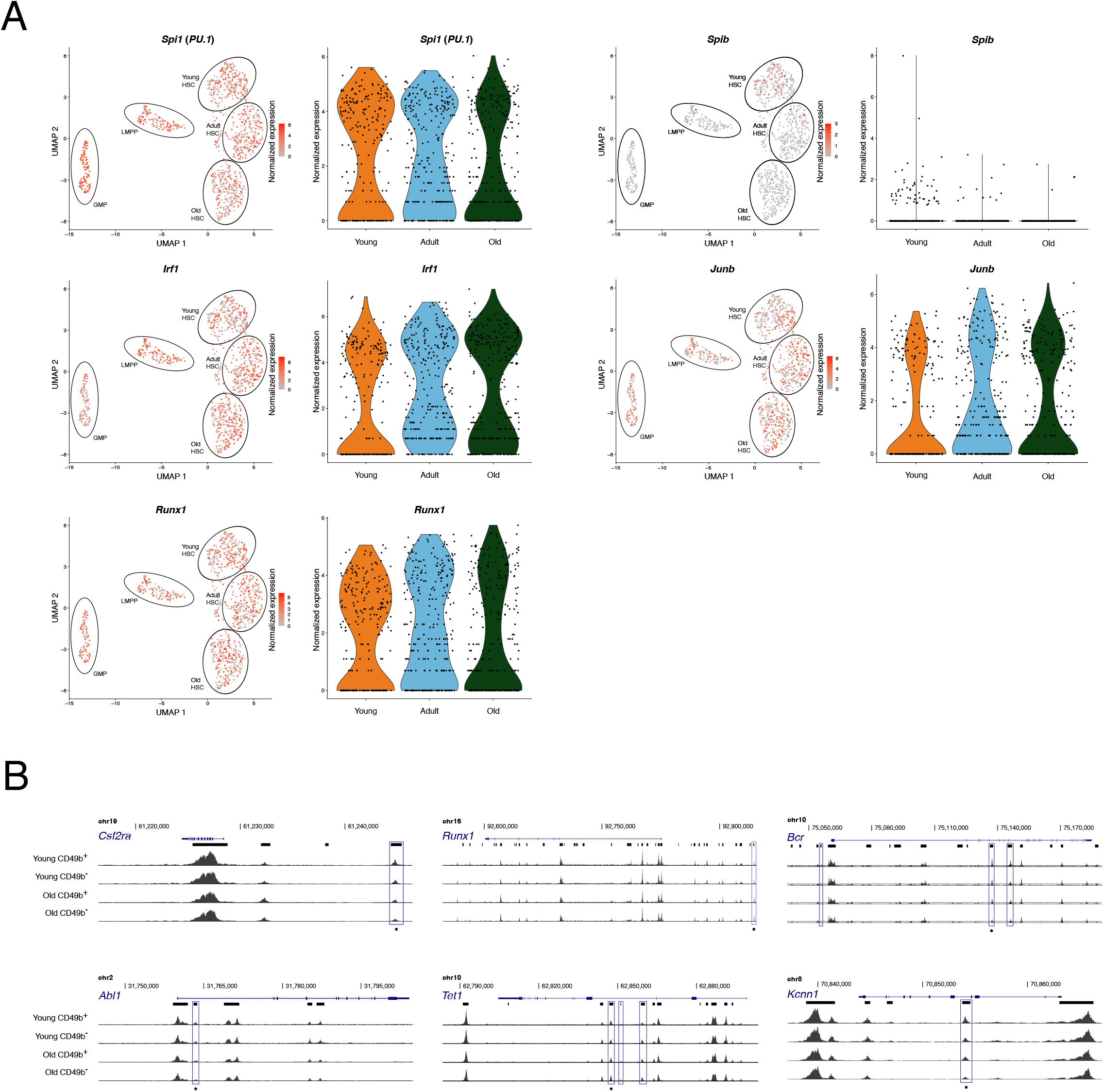
Gene expression and chromatin accessibility of selected gene regions associated with lineage bias differentiation. **a**, Expressions of *Spi1*, *Spib*, *Irf1, Junb*, and *Runx1* across stem-and progenitor cells from young, adult, and old mice. Normalized expression levels are visualized as color intensity in the UMAP and violin plots for all genes. For *Spib*, the UMAP color scale was cut-off at a maximum normalized expression of 3 to improve visibility. **b**, UCSC browser tracks of median ATAC-seq signal for selected regions. Peaks associated with lineage bias differences (Lin DARs) are indicated with blue boxes. Boxed regions indicated with asterisks are shown in Fig. 6f.

## Notes

### Competing Interest Statement

The authors have declared no competing interest.

