## Supplementary Tables 3-5 for "Aging is associated with functional and molecular changes in distinct hematopoietic stem cell subsets"

**Supplementary Table 3. Antibody list**

| Antibody | Clone | Supplier | Catalog number |
|---|---|---|---|
| **Hematopoietic stem- and progenitor cell markers** | | | |
| Anti-mouse Sca-1 (Ly-6A/E) BV605 | D7 | BioLegend | 108134 |
| Anti-mouse Sca-1 (Ly-6A/E) BV650 | D7 | BD Biosciences | 740450 |
| Anti-mouse CD117 (c-kit) APC-eF780 | 2B8 | ThermoFisher Scientific | 47-1171-82 |
| Anti-mouse CD48 AF700 | HM48-1 | BioLegend | 103426 |
| Anti-mouse CD48 APC | HM48-1 | BioLegend | 103412 |
| Anti-mouse CD150 (SLAM) BV785 | TC15-12F12.2 | BioLegend | 115937 |
| Anti-mouse CD150 (SLAM) PE-Cy7 | TC15-12F12.2 | BioLegend | 115914 |
| Anti-mouse CD34 FITC | RAM34 | ThermoFisher Scientific | 11-0341-85 |
| Anti-mouse CD34 AF647 | RAM34 | BD Biosciences | 560230 |
| Anti-mouse CD49b PE-Cy7 | HMα2 | BioLegend | 103518 |
| Anti-mouse CD49b AF647 | HMα2 | BioLegend | 103511 |
| Anti-mouse CD49b BV711 | HMα2 | BD Biosciences | 740704 |
| Anti-mouse CD105 (Endoglin) BV650 | MJ7/18 | BD Biosciences | 740609 |
| Anti-mouse CD127 (IL7-Ra) BV711 | A7R34 | BioLegend | 135035 |
| Anti-mouse CD135 (Flt-3) PE | A2F10 | BioLegend | 135306 |
| Anti-mouse CD135 (Flt-3) BV421 | A2F10.1 | BD Biosciences | 562898 |
| Anti-mouse CD16/32 AF700 | 93 | ThermoFisher Scientific | 56-0161-82 |
| Anti-mouse CD16/32 BUV737 | AB93 | BD Biosciences | 751697 |
| Anti-mouse CD16/32 (Fc-block) Purified | 2.4G2 | BD Biosciences | 553142 |
| **Donor and recipient cell markers** | | | |
| Anti-mouse CD45.1 (Ly5.1) BV421 | A20 | BD Biosciences | 563983 |
| Anti-mouse CD45.1 (Ly5.1) BUV395 | A20 | BD Biosciences | 565212 |
| Anti-mouse CD45.2 (Ly5.2) PE | 104 | BioLegend | 109808 |
| Anti-mouse CD45.2 (Ly5.2) APC-Fire750 | 104 | BioLegend | 109852 |
| **Mature hematopoietic cell markers** | | | |
| Anti-mouse CD11b (Mac-1) BUV395 | M1/70 | BD Biosciences | 563553 |
| Anti-mouse CD11b (Mac-1) BUV737 | M1/70 | BD Biosciences | 564443 |
| Anti-mouse/Human CD11b (Mac-1) BV510 | M1/70 | BioLegend | 101263 |
| Anti-mouse CD3e AF700 | 500A2 | BD Biosciences | 557984 |
| Anti-mouse CD3e BV510 | 145-2C11 | BioLegend | 100353 |
| Anti-mouse CD3e BUV395 | 1451-2C11 | BD Biosciences | 563565 |
| Anti-mouse CD4 BUV395 | RM4-5 | BD Biosciences | 740208 |
| Anti-mouse CD5 BV510 | 53-7.3 | BioLegend | 100627 |
| Anti-mouse CD5 BUV395 | 53-7.3 | BD Biosciences | 740206 |

**Supplementary Table 3. Antibody list (continued)**

| | | | |
|---|---|---|---|
| Anti-mouse CD8a BUV395 | 53-6.7 | BD Biosciences | 563786 |
| Anti-mouse Thy1.2 BV605 | 53-2.1 | BioLegend | 140318 |
| Anti-mouse CD19 PE-Cy7 | 6D5 | BioLegend | 115520 |
| Anti-mouse CD45R/B220 BUV395 | RA3-6B2 | BD Biosciences | 563793 |
| Anti-mouse CD45R/B220 BUV661 | RA3-6B2 | BD Biosciences | 565077 |
| Anti-mouse/Human CD45R/B220 PE-Dazzle 594 | RA3-6B2 | BioLegend | 103258 |
| Anti-mouse/Human CD45R/B220 BV510 | RA3-6B2 | BioLegend | 103248 |
| Anti-mouse F4/80 APC | BM8 | BioLegend | 123116 |
| Anti-mouse F4/80 APC | BM8 | ThermoFisher Scientific | 17-4801-82 |
| Anti-mouse Gr-1 BV510 | RB6-8C5 | BioLegend | 108437 |
| Anti-mouse Gr-1 BUV395 | RB6-8C5 | BD Biosciences | 563849 |
| Anti-mouse Gr-1 (Ly-6G/Ly-6C) PE | RB6-8C5 | BD Biosciences | 553128 |
| Anti-mouse NK1.1 BUV395 | PK136 | BD Biosciences | 564144 |
| Anti-mouse Ter-119 BV510 | TER-119 | BD Biosciences | 563995 |
| Anti-mouse Ter-119 BV650 | TER-119 | BD Biosciences | 747739 |
| Anti-mouse Ter-119 BUV395 | TER-119 | BD Biosciences | 563827 |
| Anti-mouse Ter-119 APC | TER-119 | Proteintech | APC-65149 |
| Anti-mouse CD41 BV786 | MWReg30 | BD Biosciences | 740903 |
| Anti-mouse CD41 PE | eBioMWReg30 | ThermoFisher Scientific | 12-0411-83 |
| **Viability, cell cycle and proliferation markers** | | | |
| 7-AAD | | BD Biosciences | 559925 |
| DAPI | | ThermoFisher Scientific | D3571 |
| Ki-67 FITC | | BD Biosciences | 556026 |
| Ki-67 PE | | BD Biosciences | 567719 |
| BrdU PE | | BD Biosciences | 556029 |

**Supplementary Table 4. Immunophenotypic definition of hematopoietic cells**

| | |
|---|---|
| **Hematopoietic stem cell compartment** | |
| $CD49b^{-}$ subset | $Lin^{-}$ $Sca\text{-}1^{+}$ $c\text{-}kit^{+}$ $CD48^{-}$ $CD34^{-}$ $CD150^{hi}$ $CD49b^{-}$ |
| $CD49b^{+}$ subset | $Lin^{-}$ $Sca\text{-}1^{+}$ $c\text{-}kit^{+}$ $CD48^{-}$ $CD34^{-}$ $CD150^{hi}$ $CD49b^{+}$ |
| $CD150^{hi}$ subset | $Lin^{-}$ $Sca\text{-}1^{+}$ $c\text{-}kit^{+}$ $CD48^{-}$ $CD34^{-}$ $CD150^{hi}$ |
| **Bone marrow stem- and progenitor cell compartment** | |
| Hematopoietic stem cell (HSC) | $Lin^{-}$ $Sca\text{-}1^{+}$ $c\text{-}kit^{+}$ $Flt\text{-}3^{-}$ $CD48^{-}$ $CD150^{+}$ or $Lin^{-}$ $Sca\text{-}1^{+}$ $c\text{-}kit^{+}$ $CD48^{-}$ $CD150^{+}$ |
| Common lymphoid progenitor (CLP) | $Lin^{-}$ $B220^{low}$ $Sca\text{-}1^{low}$ $c\text{-}kit^{low}$ $Flt\text{-}3^{hi}$ $IL\text{-}7Ra^{+}$ |
| Lymphoid-primed multipotent progenitor (LMPP) | $Lin^{-}$ $Sca\text{-}1^{+}$ $c\text{-}kit^{+}$ $Flt3^{hi}$ |
| Granulocyte-monocyte progenitor (GMP) | $Lin^{-}$ $Sca\text{-}1^{-}$ $c\text{-}kit^{+}$ $CD41^{-}$ $CD150^{-}$ $CD16/32^{+}$ |
| Megakaryocyte progenitor (MkP) | $Lin^{-}$ $Sca\text{-}1^{-}$ $c\text{-}kit^{+}$ $CD150^{+}$ $CD41^{+}$ |
| **Mature blood lineage cell compartment** | |
| Platelets | $Ter\text{-}119^{-}$ $CD41^{+}$ $CD150^{+}$ |
| Erythrocytes | $CD41^{-}$ $CD150^{-}$ $Ter\text{-}119^{+}$ |
| Myeloid cells | $CD41^{-}$ $Ter\text{-}119^{-}$ $CD3e^{-}$ $Thy1.2^{-}$ $NK1.1^{-}$ $CD49b^{-}$ $B220^{-}$ $CD19^{-}$ $CD11b^{+}$ |
| B cells | $CD41^{-}$ $Ter\text{-}119^{-}$ $CD3e^{-}$ $Thy1.2^{-}$ $NK1.1^{-}$ $CD49b^{-}$ $CD11b^{-}$ $Gr\text{-}1^{-}$ $B220^{+}$ $CD19^{+}$ |
| T cells | $CD41^{-}$ $Ter\text{-}119^{-}$ $NK1.1^{-}$ $CD49b^{-}$ $CD11b^{-}$ $Gr\text{-}1^{-}$ $B220^{-}$ $CD19^{-}$ $CD3e^{+}$ $Thy1.2^{+}$ |
| NK cells | $CD41^{-}$ $Ter\text{-}119^{-}$ $B220^{-}$ $CD19^{-}$ $CD11b^{-}$ $Gr\text{-}1^{-}$ $CD3e^{-}$ $Thy1.2^{-}$ $NK1.1^{+}$ $CD49b^{+}$ |
| **OP9 B and myeloid lineage differentiation assay** | |
| Myeloid cells | $CD11b^{+}Gr\text{-}1^{+}$ and/or $CD11b^{+}F4/80^{+}$ |
| B cells | $CD19^{+}B220^{+}$ |

**Supplementary Table 5. Culture conditions for *in vitro* assays**

| Assay type | Final concentration | Supplier | Catalog number |
|---|---|---|---|
| **OP9 B and myeloid lineage differentiation assay** | | | |
| rmSCF | 25 ng/ml | Peprotech | 250-03 |
| rhFlt3-L | 25 ng/ml | Peprotech | 300-19 |
| rhIL-7 | 20 ng/ml | Peprotech | 200-07 |
| Penicillin-Streptomycin | 1% v/v | Cytiva Hyclone | SV30010 |
| β-mercaptoethanol | 0.1mM | Merck | M6250 |
| Fetal bovine serum (FBS) | 10% v/v | Cytiva Hyclone | SH30071 |
| Opti-MEM with GlutaMAX | | Gibco | 51985-026 |
| **Cell division assay** | | | |
| rmSCF | 50 ng/ml | Peprotech | 250-03 |
| rhFlt3-L | 50 ng/ml | Peprotech | 300-19 |
| rhTpo | 50 ng/ml | Peprotech | 300-18 |
| rmIL-3 | 20 ng/ml | Peprotech | 213-13 |
| β-mercaptoethanol | 0.1mM | Merck | M6250 |
| Fetal bovine serum (FBS) | 10% v/v | Cytiva Hyclone | SH30071 |
| x-vivo15 with Gentamicin and L-glutamine | | Lonza | BE02-060F |
| **Megakaryocyte differentiation assay** | | | |
| rmSCF | 50 ng/ml | Peprotech | 250-03 |
| rmFlt3-L | 50 ng/ml | Peprotech | 300-19 |
| rhTpo | 10 ng/ml | Peprotech | 300-18 |
| rmIL-3 | 20 ng/ml | Peprotech | 213-13 |
| β-mercaptoethanol | 0.1mM | Merck | M6250 |
| Fetal bovine serum (FBS) | 10% v/v | Cytiva Hyclone | SH30071 |
| BIT 9500 | 20% v/v | Stem Cell Technologies | 9500 |
| x-vivo15 with Gentamicin and L-glutamine | | Lonza | BE02-060F |
